# Antibody-Recruiting Protein-Catalyzed Capture Agents to Combat Antibiotic-Resistant Bacteria

**DOI:** 10.1101/822346

**Authors:** Matthew N. Idso, Ajay Suresh Akhade, Mario L. Arrieta-Ortiz, Bert T. Lai, Vivek Srinivas, James P. Hopkins, Ana Oliveira Gomes, Naeha Subramanian, Nitin Baliga, James R. Heath

## Abstract

Antibiotic resistant infections are projected to cause over 10 million deaths by 2050, yet the development of new antibiotics has slowed. This points to an urgent need for methodologies for the rapid development of antibiotics against emerging drug resistant pathogens. We report on a generalizable combined computational and synthetic approach, called antibody-recruiting protein-catalyzed capture agents (AR-PCCs), to address this challenge. We applied the combinatorial PCC technology to identify macrocyclic peptide ligands against highly conserved surface protein epitopes of carbapenem-resistant *Klebsiella pneumoniae*, an opportunistic gram-negative pathogen with drug resistant strains. Multi-omic data combined with bioinformatic analyses identified epitopes of the highly expressed MrkA surface protein of *K. pneumoniae* for targeting in PCC screens. The top-performing ligand exhibited high-affinity (EC_50_∼50 nM) to full-length MrkA, and selectively bound to MrkA-expressing *K. pneumoniae*, but not to other pathogenic bacterial species. AR-PCCs conjugated with immunogens promoted antibody recruitment to *K. pneumoniae*, leading to phagocytosis and phagocytic killing by macrophages. The rapid development of this highly targeted antibiotic implies that the integrated computational and synthetic toolkit described here can be used for the accelerated production of antibiotics against drug resistant bacteria.

## Introduction

The emergence of antibiotic-resistant bacteria represents a major threat to human health,(1–4) with both increasing mortality rates and costs of care.(5–8) One example is carbapenem-resistant *Klebsiella pneumoniae* (*K. pneumoniae*) strains that harbor resistance against many or all antibiotics,(1,6,9,10) cause high rates of hospital-acquired infections(11) with associated high mortality rates.(12) Compounding the general problem of antibiotic resistance are challenges in developing and approving new antibiotics.(13) This has spurred research into understanding resistance mechanisms,(14,15) host defences,(16–18) diagnostics(19) and antibiotic generation.(20,21) Ultimately, without a rapid and perhaps general method to develop new targeted antibiotics, therapies might relapse towards those of the pre-antibiotic era.(4,13)

Immunomodulation is an emerging strategy to treat bacterial infections.(22,23) The guiding principle is to employ an immunogenic agent to elicit a targeted immune response against a particular bacterial pathogen. The archetypical immune agent is an antibody and, while many therapeutic antibodies simply neutralize pathogens by direct interaction, a few operate by enhancing immune responses against the antibody target.(22) Examples include antibodies in the Zmapp cocktail developed against the Ebola virus,(24) *Staphylococcus aureus*,(25) and *K. pneumoniae*.(26,27) While effective, antibodies can be challenging to produce and globally distribute at scale.(28,29) Alternative compelling strategies include employing synthetic molecules that bind bacterial surface proteins(30,31) or peptidoglycans,(30,32) and present haptens so as to recruit the native immune system to promote bacterial clearance. Other synthetic approaches include metabolically incorporating non-native haptens into bacterial surface components.(33,34) While inexpensive and scalable, these technologies can be challenging to adapt to different pathogens, or they can be non-selective, raising concerns about deleterious off-target effects.

Guided by these approaches, we envisioned a new class of highly targeted antibiotics called antibody-recruiting protein-catalyzed capture agents (AR-PCC) that could be rapidly developed against a specified drug-resistant bacterium (Figure 1). AR-PCCs consist of two molecular motifs. The first is a macrocyclic polypeptide ligand (the PCC) developed against a designated epitope of a specific bacterial surface protein. The second is an immunogenic antibody-recruiting (AR) label on the PCC that promotes pathogen phagocytosis by innate immune cells (Figure 1). To generate the AR-PCC ligand, we employed the recently reviewed,(35) all synthetic epitope-targeted protein-catalyzed capture agent (PCC) method,(36–39) coupled with a bioinformatics approach to identify epitopes for targeting. By targeting highly exposed, antigenic epitopes of the Type 3 Fimbrial Shaft (MrkA) surface protein of *K. pneumoniae*, we developed a small macrocyclic peptide binder in a single generation screen. That binder exhibits high affinity for the MrkA protein, high selectivity for carbapenem-resistant *K. pneumoniae*, and, when labeled with the AR tag, selectively promotes macrophage-mediated phagocytosis of the pathogen. This work demonstrates that AR-PCCs can be used to target multi-drug resistant *K. pneumoniae*, and that the basic technology might provide a route towards drugging “undruggable” pathogenic bacteria.

**Figure 1.**
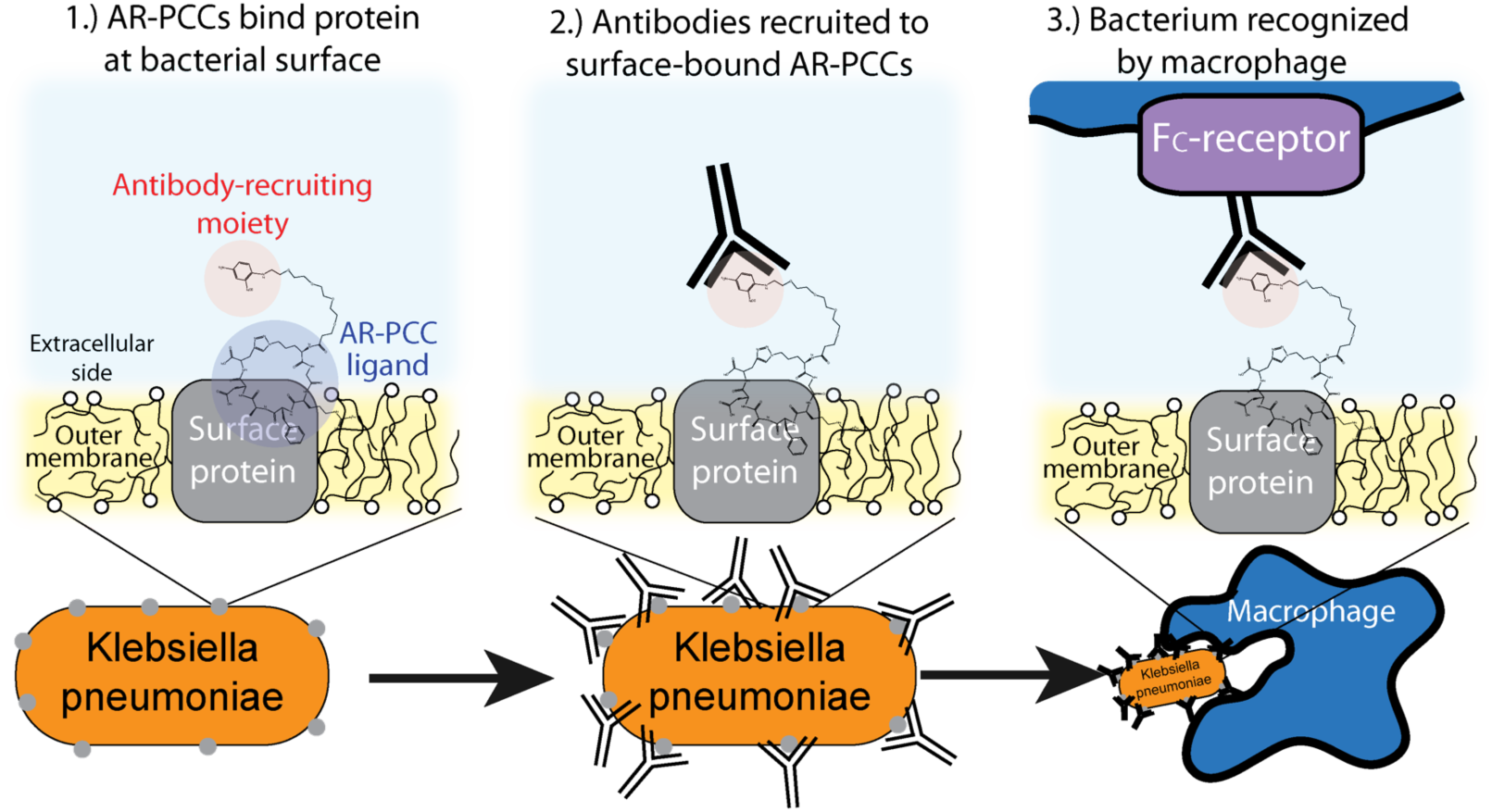
Schematic cartoon that depicts the mode-of-action of AR-PCCs against antibiotic-resistant *Klebsiella pneumoniae*. AR-PCC molecules (1) bind to a surface protein on a *K. pneumoniae* bacterium, then (2) recruit antibodies that opsonize the bacterium, which leads to (3) recognition and phagocytosis of the bacterium by immune cells (e.g., macrophages).

## Results and Discussion

### Multi-omic analyses to select target a protein on K. pneumoniae

Here we describe the algorithm used to identify protein targets, and epitopes on those targets, for drugging *K. pneumoniae* using AR-PCCs. Traditional drugging strategies tend to rely on disrupting the function of, for example, an enzyme by competing for occupancy within a strategic hydrophobic binding pocket. Our requirements are very different. Instead, favorable aspects of target proteins are high expression levels on only the pathogen of interest, plus localization of that protein to the outer membrane or extracellular space of the pathogen. Further, once such a target protein is identified, there are additional considerations regarding which epitopes of that protein present the greatest opportunities for exploiting AR-PCCs. The flow diagram in Figure 2A delineates the strategy for identifying MrkA as an ideal target protein. Protein expression levels can vary across environments and growth phases, so we analyzed the transcriptional data reported by Guilhen et al(40) to identify proteins with consistently high transcript levels across three major life phases of *K. pneumoniae*: exponential phase, stationary phase, and biofilms (including detached cells) (see Methods). Briefly, we focused on the top 10% highly-expressed genes across these life phases (515 genes out of 5146 genes in the transcriptomics dataset). Subsequent cross-referencing of these 515 genes with proteomics-derived information about the localization of *K. pneumoniae* proteins(41–43) elucidated 13 highly expressed genes that encode proteins localized to the outer membrane or extracellular space. A literature search (see Methods section for specific references) then narrowed the selection by prioritizing essential virulence- and pathogenicity-related genes as well as protein orientation in the outer membrane, to five proteins: (i) FhuA, a siderophore, (ii) Lpp, a lipoprotein, (iii) Pal, a peptidoglycan-associated lipoprotein, (iv) NlpD, a lipoprotein, (v) MrkA, a subunit of the type 3 fimbriae. Ultimately, MrkA was chosen as a target due to its key roles in infection and persistence,(44) its presence in the majority of sequenced *K. pneumoniae* strains,(45–47) and, critically, its location in fimbrial rods. These rods are large extracellular structures (0.5-2 μm long, 4-to-5 nm in diameter) (48,49) that are each comprised of up to 1,000s of MrkA copies.(50)

**Figure 2.**
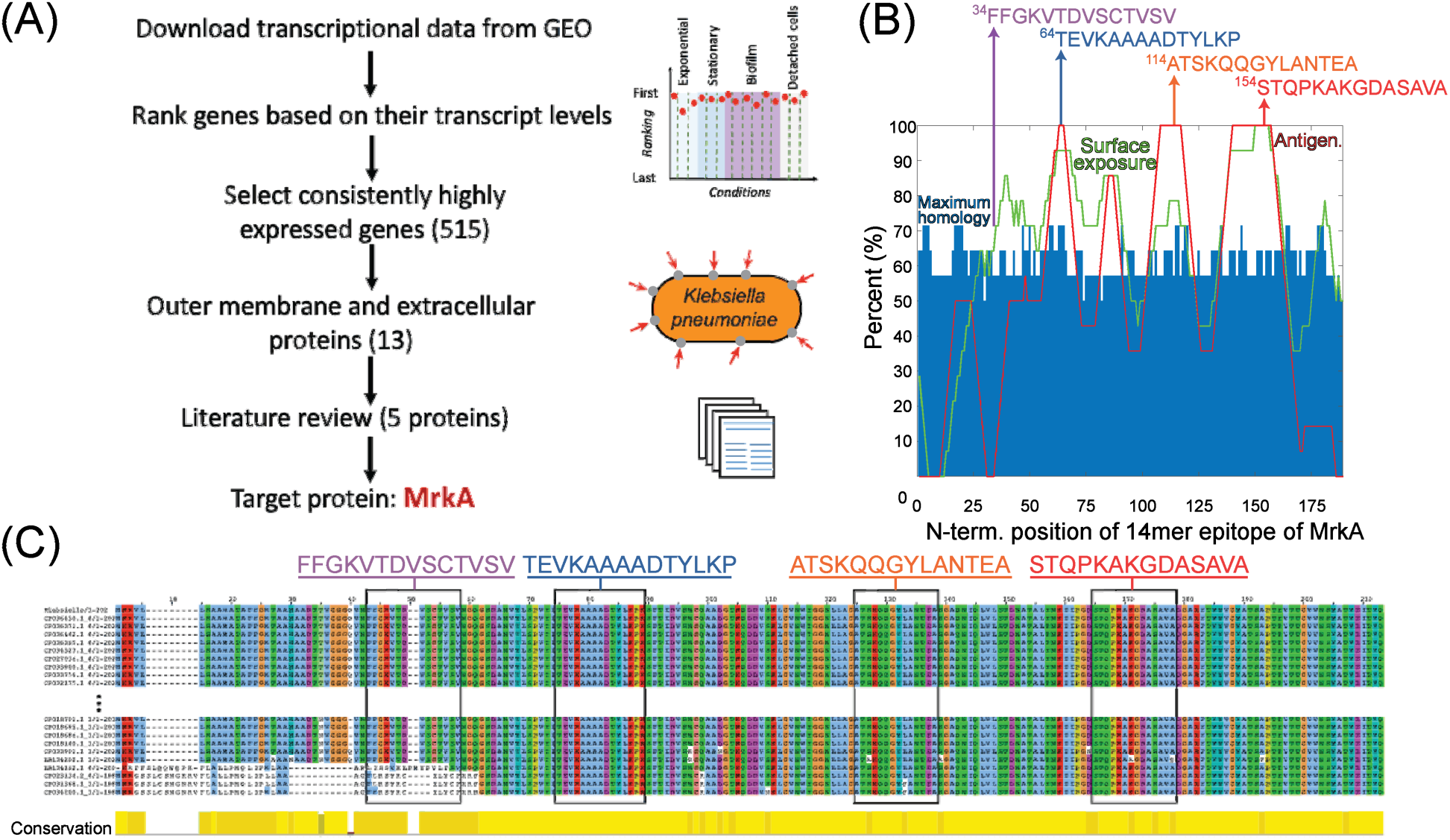
**(A)** Flow diagram illustrating the multi-omic-driven approach for selecting a protein target on *K. pneumoniae* for AR-PCCs: analysis of transcriptomic and proteomic data identified highly expressed surface proteins of *K. pneumoniae*. Among those proteins, MrkA was targeted due to its association with virulence and extracellular localization (literature review). **(B)** Plot of predicted surface-exposure, antigenicity (“Antigen.”), and maximum homology with the human proteome for all 14-residue epitopes of MrkA, as derived from bioinformatics tools (NetSurfP-2.0,(51) BepiPred 2.0,(52) and BLAST, using parameters defined in ref. (53)). Arrows indicate the locations of four 14-residue epitopes with high-exposure and low-homology that were selected as targets. The sequences of these epitopes are shown above the plot. (**C**) Alignment analyses of MrkA protein sequences from 380 *K. pneumoniae* strains in which target epitopes are shown in rectangles. Alignments were calculated by using Multiple Alignment using Fast Fourier Transform (MAFFT) and visualized in Jalview.(54,55) Only the first and last 10 sequences of the total 380 sequences are shown here.

The selection of target epitopes on MrkA is illustrated in Figure 2B. Selection of surface exposed epitopes would be aided by protein structure, but no published structures exist for MrkA. As a surrogate, we employed bioinformatics tools to survey MrkA for epitopes with high surface exposures and low homology with the human proteome. B-cell antigenicity was also mapped, but wasn’t a critical selection consideration.

The surface exposure, antigenicity, and homology of all 14-residue epitopes on the MrkA sequence were predicted and superimposed in Figure 2B. Surface-exposure and antigenicity were calculated by averaging the predicted values assigned to each residue in the full-length MrkA sequence (by BepiPred-2.0 or NetSurfP-2.0)(51,52) over the entire 14-residue epitope, while homology was predicted by comparing each 14-residue epitope with the human proteome using BlastP2.0 with parameters defined in the heuristic string method by Berglund et al.(53) This homology search produced many partial matches per epitope, and the “maximum homology” value plotted in Figure 2B represents the percentage overlap with the best match. Predicted and averaged values from these analyses are tabulated in Table S1. The plot in Figure 2B reveals several regions containing epitopes of high (>60%) predicted surface exposure (Figure 2B, green line) and relatively low (∼50-55%) maximum homology to the human proteome (Figure 2B, blue bars). Note that epitopes in surface-exposed regions range in predicted antigenicity from 0 to 100%. Four epitopes, indicated by arrows in Figure 2B, were selected based on their predicted high surface exposures (>60%), limited maximum homology (57-71%), and broad range of antigenicity values. The sequences of these candidate epitopes are shown in Figure 2B. Importantly, the MrkA protein and candidate epitopes are highly conserved among *K. pneumoniae* strains (Figure 2C). In fact, the MrkA sequence (with 202 amino acids) is 95% conserved across 380 analyzed *K. pneumoniae* strains.(54,55)

### Epitope-targeted ligands against the MrkA protein

AR-PCC ligands against the four selected epitopes were identified from a combinatorial library of macrocyclic peptides by using the epitope-targeted PCC method.(36–39) This method exploits non-catalyzed click chemistry via an *in sit*u click screen. For the screen, an alkyne-presenting, one-bead one-compound (OBOC) combinatorial library of approximately 1M peptide macrocycles with a 5-residue variable region is screened against synthetic variants of the epitopes (SynEps). Each SynEp is designed with a biotin assay handle and a strategically incorporated azide click handle (Supplementary Figures S1 and S2 contain the structures and characterization data for all synthetic epitopes).(36) The concept behind the screen is that select OBOC library elements will bind to a SynEp in just the right orientation so as to promote the azide-acetylene click reaction, thus covalently linking the SynEp to the bead. This product can be detected using the biotin assay handle on the SynEp, coupled with enzymatic amplification, to add color to the hit bead. Prior to screening SynEps, the library is cleared of beads that bound the detection antibody, streptavidin-alkaline phosphatase (SAv-AP). Hit beads are separated, and the hit candidate peptides are cleaved and sequenced using tandem mass spectrometry.

For this work, we performed a single screen of the OBOC library against all four target MrkA SynEps. The screen yielded 26 hits (Figure S3, SI) that were sequenced, scaled up, and tested for binding to full-length recombinant MrkA by single-point sandwich Enzyme-linked immunoassays (ELISAs) (Figure S4, SI). The top-performing ligand had the sequence cy(LLFFF) (Figure 3A inset), (where “cy” represents azidolysine-propargylglycine cyclization and the letters represent single-letter amino acid codes). This ligand exhibited an EC_50_ value of 50 nM to full-length recombinant MrkA protein (Figure 3A). Additional ligands, but with lower affinities to MrkA, were also discovered, with sequences cy(TTFFF), cy(YRHLG) and cy(GVHRL). Based upon chemical homology, cy(TTFFF) likely binds the same epitope as cy(LLFFF), but cy(YRHLG) and cy(GVHRL) presumably bind a different one.

**Figure 3.**
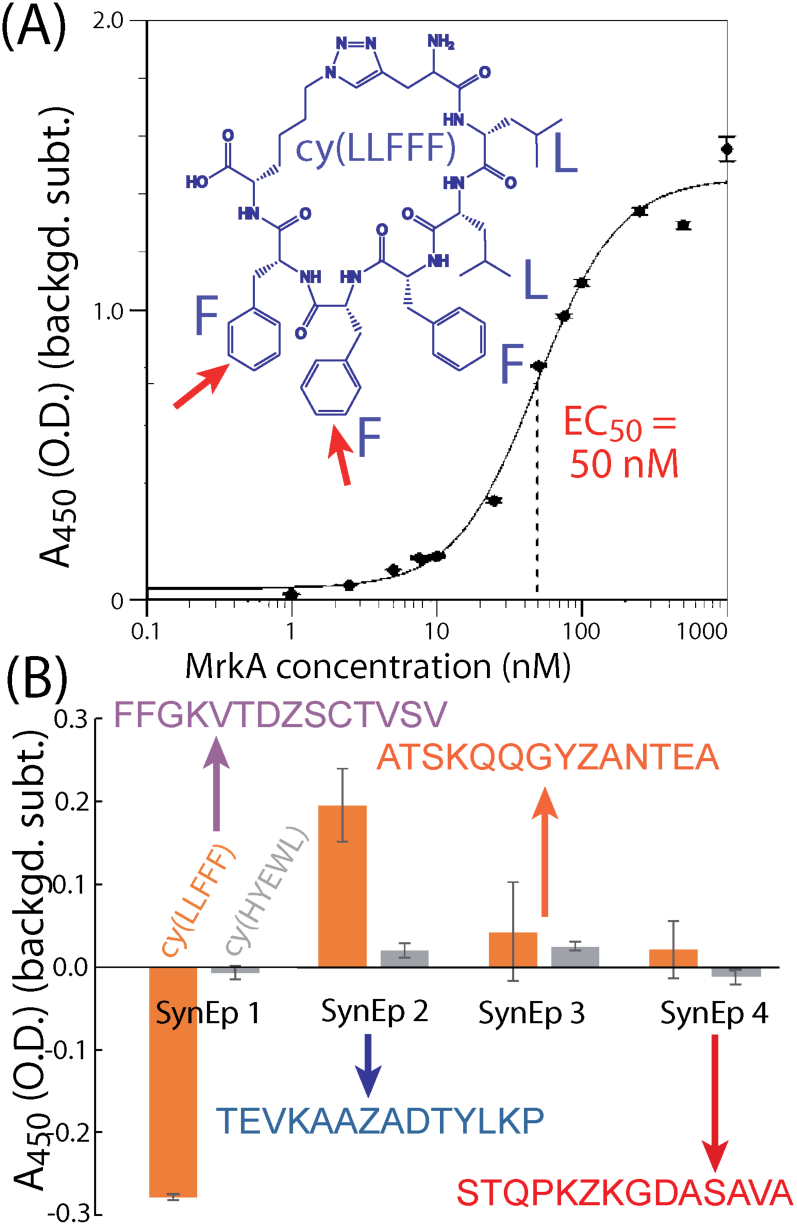
Sandwich enzyme-linked immunoassays (ELISA) reveal that the lead AR-PCC ligand cy(LLFFF) (A) binds to full-length MrkA with an EC_50_ value of 50 nM and (B) selectively binds synthetic epitope 2 versus other targeted epitopes (sequences given). The cy(HYEWL) ligand a negative control. All measurements were performed in triplicate, and all signals were background-corrected by subtracting the absorbance from otherwise identically prepared wells, except with biotin-peg6 conjugated. Error bars reflect standard deviations.

We next identified the particular epitope target to which cy(LLFFF) binds. We performed a sandwich ELISA in which the biotinylated SynEps were immobilized. The lead ligand cy(LLFFF), conjugated to a 2,4-dinitrophenyl (DNP) moiety, was titrated at a 500 nM concentration, and anti-DNP was used as a detection antibody. The results in Figure 3B show the highest signals from SynEp 2 (TEVKAAZADTYLKP). A background level for this assay was established using a 6-mer polyethylene glycol (biotin-PEG_6_). Negative signals observed for the case of SynEp 1 indicate that this SynEp blocks non-specific binding of cy(LLFFF) or of anti-DNP more than biotin-peg6. Overall, these findings suggest that cy(LLFFF) binds the epitope TEVKAAAADTYLKP in native MrkA. This epitope is 100% conserved across the 380 *K. pneumoniae* isolates analyzed (Figure 3C), suggesting that cy(LLFFF) would bind the entire cohort of these *K. pneumoniae* strains.

To further interrogate cy(LLFFF) binding, an alanine scan was used to establish which residues contribute most to binding. In this assay, a sandwich ELISA is used to quantify MrkA-binding of several cy(LLFFF) analogues in which one residue is substituted with an alanine. The ELISA results in Figure 4 show lower signal for every alanine-substituted cy(LLFFF) analogue compared to unmodified cy(LLFFF), establishing that the native compound has the highest affinity. Modestly lower binding is observed for cy(ALFFF), cy(LAFFF), and cy(LLAFF), but binding is substantially abrogated for analogues with alanine substitutions at the N-terminal diphenylalanine motif. Thus, these two N-terminal phenylalanine residues (Figure 3A, red arrows in inset) play critical roles in MrkA binding, and are targets for modifications to improve binding. Another important detail of these alanine scan results is a strong dependence of MrkA binding on the residue position, independent of residue type. For example, cy(LLAFF) and cy(LLFFA) have dramatically different binding affinities, despite having the same phenylalanine-to-alanine substitution. The indicates that MrkA-binding is sequence dependent, suggesting selectivity of cy(LLFFF) for MrkA, likely at the TEVKAAAADTYLKP epitope.

**Figure 4.**
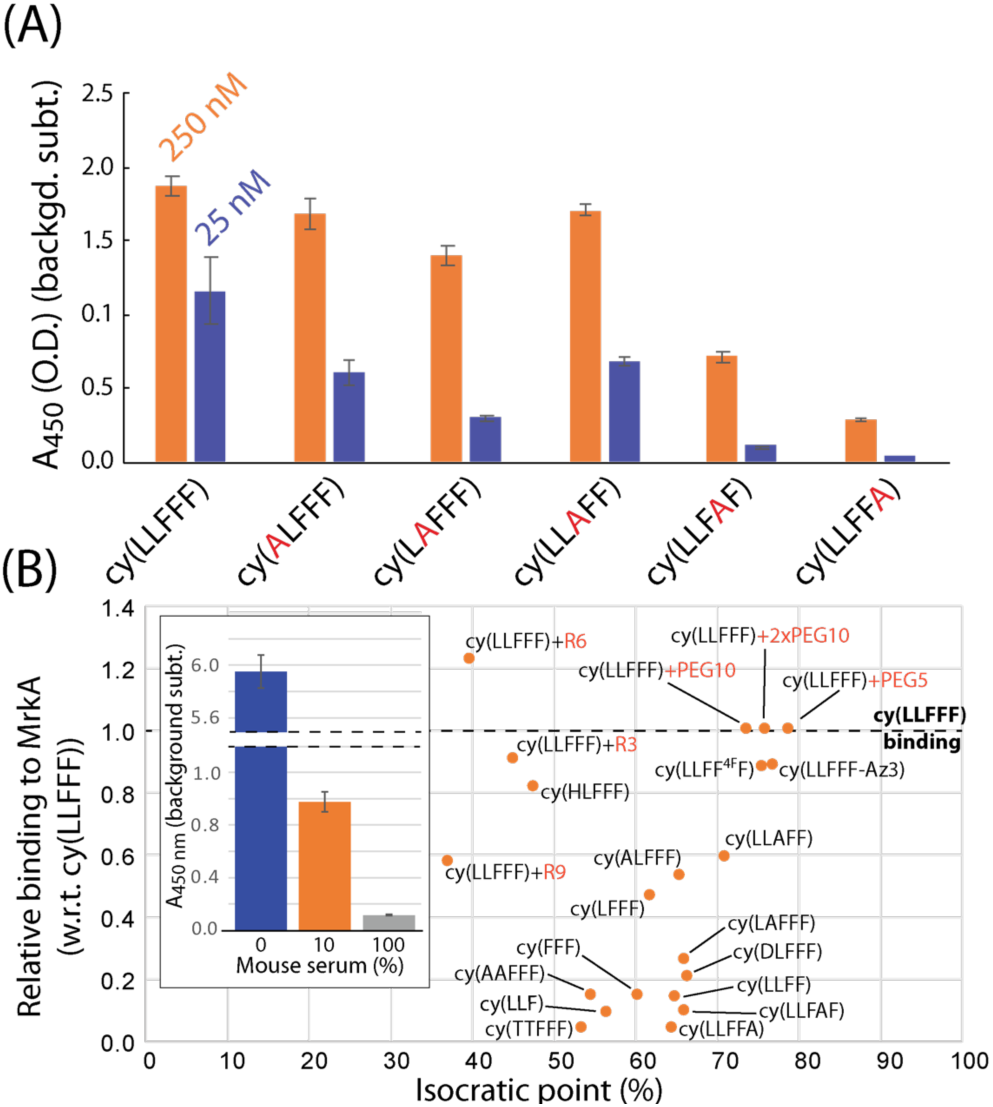
**(A)** Sandwich ELISA results of an alanine scan of cy(LLFFF), which interrogates the affinity of alanine-substituted cy(LLFFF) analogues to full-length MrkA protein. **(B)** Plot of the isocratic point versus affinity to MrkA, relative to cy(LLFFF), for various cy(LLFFF) analogues, in which residues were substituted or N-terminal tags were appended. All modifications reduced isocratic point but also reduced binding, with the exception an N-terminal hexa-arginine tag (“cy(LLFFF)+R6”). This compound also showed low yet non-zero binding to MrkA in 10 and 100% mouse sera (inset). The relative affinities plotted in (B) were calculated by taking the ratio of signals from the analogue to those of a cy(LLFFF) reference on the same ELISA plate. All ELISA data are background corrected to signal from an otherwise identical measurement except with biotin-peg6 conjugated instead of the PCC. Error bars are standard deviations.

### Synthetic modifications to optimize AR-PCC pharmacokinetics and avidity

The efficacy of AR-PCCs *in vivo* will not only depend on target binding, but also pharmacokinetic (PK) properties, of which clearance pathway is a critical parameter. Separate mouse studies in our labs indicate that PCCs with isocratic points (IPs) above 35% predominantly clear via the liver (i.e., hepatic clearance), while more hydrophilic compounds with lower isocratic points clear by the kidneys (i.e., renal clearance).(56) Thus, the highly hydrophobic cy(LLFFF) ligand would likely exhibit hepatic clearance. To afford greater control over PK properties, we explored synthetic modifications to improve the hydrophilicity of cy(LLFFF), to favor renal clearance, while retaining the desired avidity characteristics. Modifications include single- and double-residue substitutions, residue removal, and the addition of non-ionic PEG and charged polyarginine tags (R*i*, where *i* represents the number of arginine). Results of this optimization are shown in Figure 4B, which plots IP versus the affinity of the compound to MrkA, relative to cy(LLFFF). The same results are also tabulated in Table S2.

Residue substitutions and removals categorically reduced affinity to MrkA, indicating that the structure of cy(LLFFF) compound is somewhat optimized to bind the TEVKAAADTYLKP epitope. These residue substitutions and removals provided moderate (up to ∼20%) reductions in IP versus cy(LLFFF). By comparison, the addition of polyarginine tags reduced the IP up to ∼40% without substantial loss in MrkA avidity (three leftmost data points in Figure 4B). In fact, cy(LLFFF) with a hexa-arginine tag shows a 22% stronger affinity to MrkA over cy(LLFFF) and had an isocratic point of near 40%—an improvement of ∼35% over unmodified cy(LLFFF). This compound also binds MrkA in mouse sera (Fig 4B inset), making this molecule promising for *in vivo* translation.

### AR-PCC binding to multidrug resistant *Klebsiella pneumoniae* surfaces

While cy(LLFFF) exhibits high affinity towards MrkA in solution, the context of MrkA is much different in native fimbriae where MrkA proteins oligomerize into a helix-like structure.(49,57) The binding of cy(LLFFF) to *K. pneumoniae* cells was tested using cell-based ELISAs (Figure 5A). For these assays, *K. pneumoniae* cells were cultivated in glycerol-casamino acid broth to induce the expression of type 3 fimbria,(48) which are predominantly composed of MrkA, and MrkA expression was verified by using Western Blot (Figure S5). These cells were exposed to biotin-PCC ligands, then SAv-horseradish peroxidase (SAv-HRP), and then developed to produce a colorometric signal that correlates with SAv-HRP binding. The results in Figure 5B show a strong signal from the two strains of *K. pneumoniae* (BAA 1705 and BAA 2146) but only weak signal from identically prepared, but non-MrkA-expressing, *Escherichia coli* (*E. coli*) or *Salmonella typhimurium* (*S. typhimurium*) cells. When the same bacterial strains were exposed instead to biotinylated cy(HNGPT), a non-MrkA binding control ligand, little binding was observed, with signal comparable to untreated (Figure 5B) and secondary antibody controls (Figure S6). The lack of binding by cy(HNGPT), and comparative strong binding of cy(LLFFF) to *K. pneumoniae* cells demonstrate the excellent binding selectivity of biotinylated cy(LLFFF) for MrkA-expressing *K. pneumoniae* cells versus non-target bacteria. Importantly, though not explicitly tested here, *E. coli* and *S. typhimurium* are known to express fimbria under growth conditions similar to those used here,(58,59) suggesting selectivity of cy(LLFFF) to type 3 fimbriae of *K. pneumoniae* as well. Moreover, salt aggregation tests reveal that the cell surfaces of the *K. pneumoniae* strains used here have similar or lower hydrophobicities than *E. coli* or *S. typhimurium* (Figures S7 and S8), indicating that binding selectivity cannot be explained simply by hydrophobicity. These binding results also indicate that labeled PCCs can be utilized to recruit biomacromolecules (i.e., SAv-HRP) selectively to the targeted *K. pneumoniae* cells.

**Figure 5.**
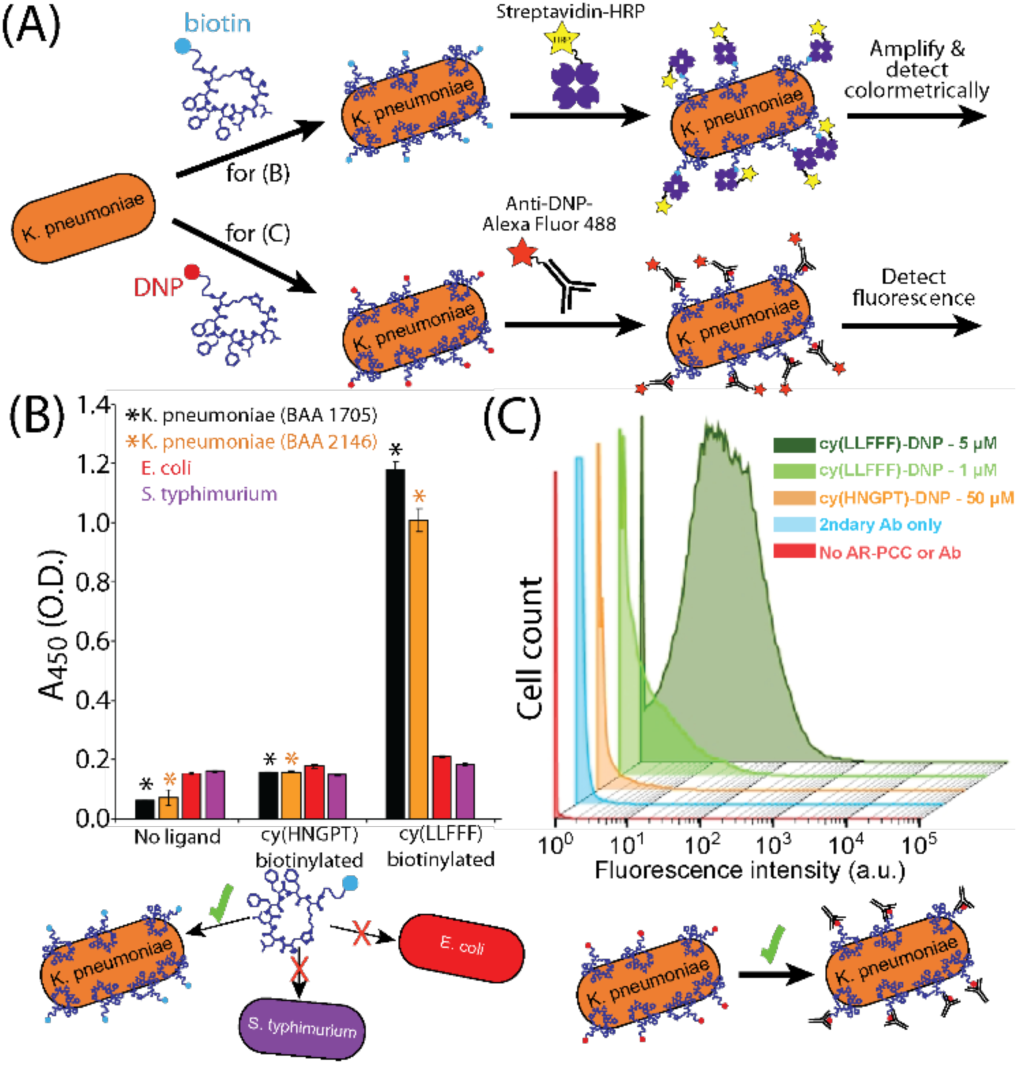
**(A)** Schematic illustration of the opsonization assays, in which *K. pneumoniae* was incubated with PCCs conjugated with either biotin or DNP, opsonized by corresponding antibody, and then detected. **(B)** Cell-binding ELISA results shows that biotinylated cy(LLFFF) at 50 μM binds two strains of MrkA-expressing *K. pneumoniae* cells with excellent selectivity versus *E. coli* and *S. typhimurium*. By comparison a biotinylated dummy ligand cy(HNGPT) shows no or minimal binding above baseline. Asterisks (*) indicate ELISA data for target *K. pneumoniae* strains, and error bars reflect standard deviations. **(C)** Flow cytometry results for *K. pneumoniae* cells (strain BAA 2146) opsonized by DNP-conjugated PCCs and fluorescently-tagged anti-DNP antibodies. Cells exposed to cy(LLFFF)-DNP, at either 1 or 5 μM concentrations, plus secondary antibody (light and dark green) showed much higher fluorescence than cells incubated with 50 μM of cy(HGNPT)-DNP conjugate plus secondary antibody (orange) and secondary antibody only (teal). Untreated cells showed the lowest fluorescence (red). Cytometry measurements were gated for single cells, and each distribution comprises >18,000 cells.

### AR-PCC-driven opsonization of *K. pneumoniae*

We next sought to use cy(LLFFF) to promote opsonization by replacing the biotin tag with the immunogen 2,4-dinitrophenyl (DNP) to make the AR-PCC. DNP is an agonist for 1% of endogenous human antibodies,(60) and so once the AR-PCC binds to the *K. pneumoniae* surface, we hypothesized that the AR handle should recruit antibodies to the pathogen. PCCs were tagged with a DNP group via conjugation of a DNP-modified lysine residue, in which a DNP moiety is covalently attached at the terminal sidechain amine. Both DNP-conjugated cy(LLFFF) and cy(HNGPT) PCCs showed absorbances at 360 nm and 420 nm (Figure S9) that are characteristic of DNP-modified lysine(34) and indicate successful labeling of the PCC.

Flow cytometry was used to determine the extent to which cy(LLFFF)-DNP conjugates promote the opsonization of resistant *K. pneumoniae* by anti-DNP antibodies. *K. pneumoniae* cells (strain BAA 2146) were first exposed to cy(LLFFF)-DNP and then to Alexafluor 488 fluorophore-labeled anti-DNP antibodies (Figure 5A), so fluorescence served as a proxy for opsonization. The cytometry data in Figure 5C show weak fluorescence (predominantly <0.2×10^1^ intensity) *K. pneumoniae* cells that were either unstained or incubated with fluorescent anti-DNP only, and slightly higher fluorescence (predominantly <1.0×10^1^ intensity) was observed for cells exposed to cy(HNGPT)-DNP at 50 μM plus anti-DNP, indicating slight cross-reactivity. Fluorescence counts for all these samples, however, are substantially less than for cells treated with much lower concentrations (1 or 5 μM) of cy(LLFFF)-DNP plus fluorescent anti-DNP (Figure 4C, light and dark green, respectively). This demonstrates that cy(LLFFF)-DNP promotes the opsonization of *K. pneumoniae* cells with anti-DNP. The same behaviors were observed on a separate resistant *K. pneumoniae* strain, BAA 1705, (Figure S10) and by using cell-based ELISA methods that were sensitive to anti-DNP binding (Figure S11). Paired with the recruitment of SAv-HRP to cells by biotin-cy(LLFFF) (Figure 5A), these results lend to the versatility of AR-PCCs to recruit diverse and specified biomolecules to bacterial surfaces.

### AR-PCC-driven phagocytosis and opsonophagocytic killing (OPK) of *K. pneumoniae*

Given that cy(LLFFF)-DNP recruits antibodies to *K. pneumoniae* surfaces, we posited that *K. pneumoniae* opsonized in this manner would be more susceptible to phagocytosis by macrophages. AR-PCC-driven phagocytosis and OPK were quantified by a gentamicin protection assay, as depicted in Figure 6A. For this assay, *K. pneumoniae* cells (strain BAA 1705) were treated with cy(LLFFF)-DNP, opsonized by anti-DNP antibodies, and then incubated with murine bone-marrow-derived macrophages (BMDMs). BMDMs were subsequently exposed to a solution containing gentamicin, to which *K. pneumoniae* is susceptible, to kill any extracellular bacteria. Phagocytosed bacteria remain viable inside BMDMs for at least 1 h, yet are rendered inviable after 24 h by OPK (Figure S12). Macrophages harvested at 1 h and 24 h were lysed and plated to generate bacterial colonies, which were enumerated to quantify phagocytosis and OPK.

**Figure 6.**
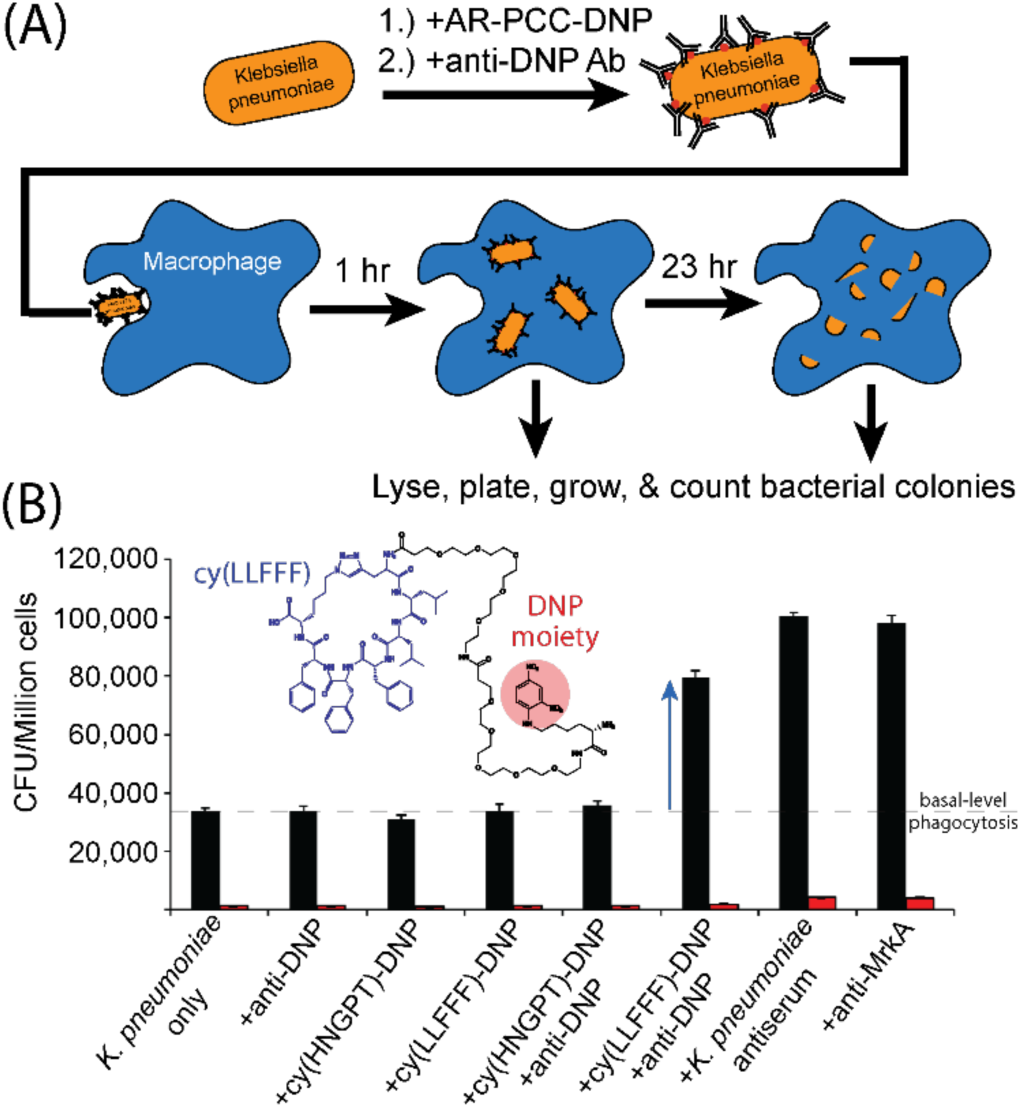
**(A)** Flow diagram that depicts the assay used to evaluate AR-PCC-driven opsonophagocytic killing (OPK) of *K. pneumoniae* cells. Briefly, *K. pneumoniae* cells (strain BAA 1705) opsonized by AR-PCCs and anti-DNP antibodies were incubated with macrophages for 1 or 24 h. Macrophages were lysed, and the lysate was plated to obtain bacterial colonies that were subsequently counted. **(B)** Plot of colony counts for various samples subjected to the OPK assay. At 1 h, all controls (the five leftmost samples) showed basal-level CFU counts at around 35,000 CFU, while *K. pneumoniae* cells treated with lead AR-PCC cy(LLFFF)-DNP plus anti-DNP antibody produced much higher 80,000 CFU (blue arrow), establishing cy(LLFFF)-DNP promotes opsonization. This level of phagocytosis is comparable to that promoted by *K. pneumoniae* antiserum and anti-MrkA antibody (at dilutions stated in the experimental section), which yielded approximately 100,000 CFU. Nearly no CFU were produced from any samples harvested at 24 h, indicating that AR-PCCs did not interfere with OPK activity. All AR-PCCs were used at a 50 µM concentration, and measurements were conducted in triplicate, with error bars here indicating standard deviations.

Phagocytosis assays were carried out using cy(LLFFF)-DNP on a strain of *K. pneumoniae* (BAA 1705) that harbored high antimicrobial resistance, including all tested carbapenems, but is susceptible to gentamicin. As shown in Figure 6B, BMDMs exposed to untreated *K. pneumoniae* cells for 1 h yielded counts of ∼35,000 CFU/million cells, establishing basal levels of phagocytosis (dashed line in Figure 6B). Similar signals were also observed in all control samples (Figure 6B, leftmost five samples) harvested at the 1 h time point. The controls were bacteria treated either with anti-DNP antibody only, cy(LLFFF)-DNP only, cy(HNGPT)-DNP only, or cy(HNGPT)-DNP plus anti-DNP. None of these treatments elicited phagocytosis above basal levels. Much greater counts of 80,000 CFU (blue arrow, Figure 6B) were observed from samples in which *K. pneumoniae* was treated with no compounds (“*K. pneumoniae* only”), cy(LLFFF)-DNP plus anti-DNP, which compared well with the OPK-promoting performances of the *K. pneumoniae* antiserum and the anti-MrkA antibody (Figure 6B, rightmost two samples). T-tests conducted on the data shown in Figure 6B establish that cy(LLFFF)-DNP plus anti-DNP significantly promoted more phagocytosis than any of the controls (p-value <0.001, Figure S13, SI). However, phagocytosis resulting from *K. pneumoniae* antiserum and anti-MrkA treatment produced statistically higher phagocytosis (p <0.001) than cy(LLFFF)-DNP plus anti-DNP. At 24 h of incubation, the cy(LLFFF) plus anti-DNP sample (and all other samples) showed little to no cell counts, indicating near complete OPK. Thus, AR-PCCs demonstrably enhance the OPK of a highly resistant *K. pneumoniae* bacterium, presumably by engaging F_C_-receptor-mediated phagocytosis pathways.

## Conclusions

We demonstrated, in vitro, a new concept for targeted antibiotics called antibody-recruiting protein-catalyzed capture agents (AR-PCCs). AR-PCCs were designed to exhibit specific *in vitro* antimicrobial activities against highly-resistant *Klebsiella pneumoniae* bacteria. AR-PCC molecules comprise of two molecular motifs: a peptide ligand that binds a specific surface protein epitope on the pathogen, and an immunogenic moiety that recruits antibodies. Combined multi-omic data and bioinformatic analyses provided an algorithm for selecting a highly abundant surface protein and epitopes on *K. pneumoniae* as targets for AR-PCCs. A single-generation PCC combinatorial screen was then used to rapidly identify macrocyclic peptide ligands to the chosen epitope targets. The lead AR-PCC ligand, cy(LLFFF), exhibited strong binding to full-length MrkA (EC_50_=50 nM) and one of the highly conserved target epitopes, as well as high specificity to MrkA-expressing *K. pneumoniae* (compared to other bacterial species that do not express MrkA). Further, the lead AR-PCC ligand conjugated with 2,4-dinitrophenyl (DNP) moieties recruited anti-DNP antibodies to *K. pneumoniae* surface, which led to increased levels of phagocytosis by macrophages and ultimately greater opsonophagocytic killing. Chemical modifications to the cy(LLFFF) suggest that the macrocycle scaffold can be optimized for desired in vivo PK characteristics. Such mouse model experiments are currently underway. While *K. pneumoniae* served as the target in this study, the approaches used here should be adaptable to other antibiotic resistant extracellular pathogens. This versatility also makes feasible the development of cocktails of AR-PCCs that simultaneously target several conserved surface epitopes on a single pathogen, to facilitate complete clearance of bacterial populations that exhibit heterogenous surface protein expression. Overall, AR-PCCs are a promising all-synthetic molecular platform that can be rapidly designed, built and deployed against resistant microbes.

## Experimental

### Gene expression analysis

Transcriptional profiles (normalized read counts) of *Klebsiella pneumoniae* strain CH1034 in stationary phase, exponential phase, and biofilms (7h, 13h and detached cells) generated by Guilhen and collaborators^36^ were downloaded from the GEO database (accession number: GSE71754).(61) The downloaded dataset included 5,146 genes and triplicates for each condition. Although additional transcriptomes are publicly available for *K. pneumoniae*, we restricted our analysis to a single dataset (with key stages of *K. pneumoniae* life cycle) due to: (i) the high number of *K. pneumoniae* strains studied by different research groups. This means available transcriptomic data involve multiple strains and have been collected on multiple platforms. (ii) the large size of *K. pneumoniae* pangenome(45) makes possible transcript levels of any given gene may be present in only some of the available transcriptional profiles. Inspired by the superior performance of rank-combined predictions that integrate multiple network inference methods/predictions (with respect to single ones(62,63)), we ranked all genes in each replicate based on their normalized read counts. Then, we computed the average ranking position of each gene along the 15 replicates. This approach identified genes with consistently high transcript levels along the sampled conditions and, for downstream analyses, we focused on the set of genes that were in the top 10% of the ranking (515 genes).

### Identification of highly expressed genes encoding outer membrane or extracellular proteins

We mined available literature to identify *K. pneumoniae* proteins that localize in the outer membrane or extracellular space.(41–43) We found 54 proteins that localize to the regions of interest. Then, we evaluated the overlap between this set of proteins and the group of 515 highly expressed genes defined in the “Gene Expression Analysis” section. To compare the two sets, we first downloaded the genome annotation of *Klebsiella pneumoniae* strain CH1034 from the NCBI website in May 2018. We used the genome annotation to convert the locus names used as gene ID in the analyzed transcriptomics data to standard gene names (e.g. CH1034_10002 corresponds to *phnV*). Finally, we manually reviewed available information(48,50,64–71) about genes present in both sets to select a target, prioritizing gene targets that yielded virulence-associated proteins that were either membrane-spanning or oriented on the extracellular side of the outer membrane.

### MrkA sequence alignment

First we performed a Blastn (megablast) in the NCBI BLAST website using as query the nucleotide sequence of *mrkA* in the *K. pneumoniae ATCC BAA-1705* strain. We restricted the search to the *K. pneumoniae* taxa and set the maximum number of allowed target sequences to 500. Other parameters were kept as default. We downloaded all hits (402). Then, we removed any hit from plasmid sequences or duplicated sequences. Nucleotide hit sequences were then translated using EMBOSS Transeq.(54) Finally, MrkA protein sequences were aligned using MAFFT and visualized in Jalview.(54,55)

### Bioinformatics analysis

Predictions for surface exposure and surface antigenicity were performed using NetsurfP 2.0(51) and Bepipred2.0(52) software (using default parameters), with the primary sequence of MrkA as an input, which assign either a value of 1 or 0 to each residue. The resulting prediction values were then averaged over 14-residue epitopes, converted to a percent value, and plotted. The uniqueness of each 14-residue epitope of MrkA among all epitopes in the human proteome was determined by performing a BlastP2.0 search with the parameters defined in the heuristic string method by Berglund et al.(53) Many matches were obtained for each epitope, and the match with greatest homology with the MrkA epitope was extracted, the corresponding homology converted to a percent value, and plotted as “maximum homology.” All results from these bioinformatics analyses are tabulated in Table S1.

### Peptide Synthesis and Purification

All peptides were synthesized by using standard Fmoc solid-phase peptide synthesis procedures, using either Rink Amide resin (Aapptech, RRZ005), Sieber Amide resin (Aapptech, RST001), or Tentagel S NH2 resin (Rapp Polymere, S30902). N-methylpyrrolidone (NMP, Alfa Aesar, 43894) was used as a solvent for all synthesis procedures, except the coupling of biotin (AK Scientific, C820) which used a 50/50 mixture of dimethylsulfoxide (DMSO, Fisher, D128-4) and NMP for solubility. All standard Fmoc-protected amino acids were purchased from ChemPep. Fmoc deprotection was achieved with 20% piperidine (Alfa Aesar, A12442) in NMP, and coupling reactions employed O-(7-Azabenzotriazol-1-yl)-N,N,N’,N’-tetramethyluronium hexafluorophosphate (HATU, Chem-Impex Int’l Inc., 12881) as an activator and N-diisopropylethylamine (DIEA, Alfa Aesar, A11801) as the base. An Aapptech Titan 357 instrument was used to couple all Fmoc-protected standard amino acids and non-natural residues with click handles, i.e. propargylglycine (Pra, ChemPep, 180710) and azidolysine (Az4, ChemPep, 101227). Fmoc-NH-peg5-CH_2_CH_2_OH (ChemPep, 280110), Fmoc-NH-peg10-CH_2_CH_2_OH (ChemPep, 280113), and Fmoc-Lys(DNP)-OH (ApexBio, A7926) were coupled overnight with molar excesses of 2, 2, and 6 for the Fmoc-protected compound, HATU, and DIEA, respectively (excesses with respect to the reactive groups on bead surface). The Cu-catalyzed azide-alkyne click reaction was conducted by incubating beads overnight with shaking in a solution of 1.5x molar excess of Cu(I) (Millipore Sigma, 818311) and 5x molar excess of L-ascorbic acid (Sigma, A0278) in 20% piperidine in NMP (excesses with respect to the reactive groups on resin surface). The resin was then washed 5×1 minute with 5-8 mL of NMP, after which extra copper was removed by incubating beads with shaking for 5 minutes in a solution of 5 w/v % DIEA, 5 v/v % Sodium diethyldithiocarbamate trihydrate (Chem-Impex) in NMP. The resin is washed extensively with NMP (3x without shaking, 5×5min or more with shaking) until the bead color turned white to light yellow and remained constant. After synthesis was complete, the resin was dried in dichloromethane (DCM, Acros Organics, 40692-0040) for at least 15 mins on a vacuum manifold. Peptides were cleaved from dried resin by mixing with 10 mL solution of Trifluoroacetic acid (TFA, Alfa Aesar, L06374):Triethylsilane (TES, Millipore Sigma, 230197) Millipore Water (H_2_O) at volumetric ratios of 95:2.5:2.5, respectively, and vigorously stirring for 2 hr. Cleavage under these acidic conditions also removed all acid-labile sidechain protecting groups. The resulting solution was added to 40 mL of diethyl ether (Acros Organics, 615080-0040) and stored overnight at −20 °C to precipitate the peptide product. The product was then pelleted by centrifugation, dried in air, and then resuspended in an aqueous solution of 30% acetonitrile (Fisher, A955-4) (aq.) with 0.1% of either TFA or formic acid (Fisher, A117-50) before purification by liquid-chromatography-mass spectrometry.

PCC compounds were purified on a Waters Autopurification system, which isolated compounds based on MS peaks corresponding to protonated [M+1H]^+^ and [M+2H]^2+^, and/or sodium adducts [M+Na]^+^ and [M+2Na]^2+^. A representative HPLC chromatogram from the Waters Autopurification system with an analytical HPLC run of the corresponding product is shown in Figure S14. The isocratic points of PCCs were determined based on the elution of the compound from a C18 prep-scale column. Synthetic epitopes were either purified on the Waters Autopurification System, or by semi-preparative HPLC and then using matrix-assisted laser desorption/ionization mass spectroscopy to identify fractions with the desired m/z ratio. The resulting peptides were lyophilized, the yield determined by mass difference or UV-visible absorbance, and resuspended at a concentration of up to 10 mM peptide in DMSO. Peptides were stored at −80°C before thawing for each use.

Standard Fmoc-protected synthesis and split-and-mix procedures were used to synthesize a one-bead one-compound library on TentaGel S NH_2_ beads with the structure NH_2_-Pra(80%)/Gly(20%)-X_1_X_2_X_3_X_4_X_5_-Az4-M-Resin, where X_i_ indicates one of 16 natural amino acids (excluding Methionine, Cysteine, Glutamine and Isoleucine). The coupling solution for the N-terminal residue included 80% Fmoc-propargylglycine-OH and 20% Fmoc-glycine-OH. The resulting library was clicked under copper-catalyzed conditions as described above to yield on each bead ∼80% of the cyclic product cy(Pra-X_1_X_2_X_3_X_4_X_5_-Az4) and 20% of the linear product Gly-X_1_X_2_X_3_X_4_X_5_-Az4, which facilitates identification by tandem mass spectroscopy. A final Pra was coupled onto the library to enable PCC combinatorial screening. The library was then incubated with a solution containing volumetric ratios of 95:2.5:2.5 of TFA:H_2_O:TES under vigorous stirring for 2 h to remove acid-labile sidechain protecting groups, and the washed 3×5 min in H_2_O, NMP, methanol (Fisher, A454-1), and then DCM.

### Combinatorial PCC screening

Combinatorial in situ click screens were performed as described previously.(38) Briefly, 500 mg (representing ∼1.4 copies of approximately 1,000,000 different compounds) of a combinatorial one-bead one-compound libraries was incubated overnight with shaking in TBS buffer (25 mM Tris HCl, 150 mM NaCl, pH 7.6). A preclear to remove beads that bound SAv-AP was performed as follows. Unless otherwise noted, all steps were performed at room temperature and all incubation and washing steps were conducted with 4 mL of the stated solution with shaking. Beads were blocked overnight by incubation in blocking buffer (TBS buffer with 1% BSA and 0.05% Tween-20, pH 7.6), rinsed with blocking buffer 3×1 min, incubated for 1h with 1:10,000 streptavidin-Alkaline phosphatase (SAv-AP) (Thermofisher Scientific, SA1008) in 5 mL of blocking buffer, and then washed with the following: 3×5 min in TBS, 3×5 min in 0.1M glycine (pH 2.8), 3×5 min in TBS buffer, 3×5 min of alkaline phosphatase (AP) buffer (100 mM Tris-HCl, 150 mM NaCl, 1 mM MgCl_2_, pH 9.0). The beads were then split and transferred into two separate petri dishes by using AP buffer, such that there was a total of 16 mL of AP buffer per dish. Separately, 10 mL of BCIP/NBT development buffer was prepared from the Promega BCIP/NBT Color Development substrate kit (Promega, S3771) by adding 66 μL of the NBT solution to 10 mL of AP buffer, mixing by hand, then adding 33 μL of the BCIP solution followed by vortexing. Four mL of the BCIP/NBT development buffer was added to each plate and then each plate was gently swirled for 30 seconds to ensure the development buffer was well-mixed. The reaction was quenched after 25 minutes by adding 4 mL of 5.0N HCl (aq.) to each plate and swirling to homogenize. The beads were transferred to a new SPPS tube by using Millipore water and were then washed copiously with Millipore water (10x without shaking, then 10×1 minute with shaking). The beads were resuspended in 0.05 N HCl (aq.), returned to a petri dish, and the purple beads removed by using a 10 □L pipette.

After all of the purple beads were removed, the in situ click screen was performed with the MrkA SynEps. The library was collected and rinsed in TBS buffer (3×5 min), incubated with a TBS buffer containing 20 μM of each purified MrkA SynEp for 6 h. The library was then subjected to the following treatments: washed with TBS buffer (10×1 min), incubated for 1 h in 7.5 M Guanidine-Hydrochloride (pH 2.0), washed with TBS (10×1 min), incubated for 2 h with blocking buffer, washed with blocking buffer (3×1 min), and incubated for 1 h with 1:10,000 SAv-AP in 5 mL of blocking buffer. The library was then washed as follows: 3×5 min in TBS, 3×5 min in 0.1M glycine (pH 2.8), 3×5 min in TBS buffer, 3×5 min of AP buffer. The library was then split into two petri dishes and developed as described above. The reaction was quenched by the addition of 4 mL of 5.0N HCl (aq.), then the beads were transferred to a new SPPS tube, washed copiously with Millipore water (10x without shaking, then 10×1 minute with shaking), resuspended in 0.05N HCl, and then returned to petri dishes. Dark and medium-dark colored beads were picked as hits and collected into Corning Costar Spin-X centrifuge tube filters (Sigma-Aldrich, CLS8170). Hit beads were then rinsed 10×30s (7,000 RPM, tabletop centrifuge), decolorized by overnight incubation in NMP, rinsed in Millipore water (10×30s), resuspended in TBS, and stored at 4 °C.

Individual beads were transferred to wells in a 96 well plate and subjected to cyanogen-bromide cleavage, prepared, and sequenced as described previously.(38,72)

### ELISA assays

ELISA assays for MrkA-binding and epitope selectivity were performed on clear NeutrAvidin Coated High Capacity Plates (ThermoFisher, 15507). The buffer used for all washes and to dissolve all compounds was TBST+0.1% BSA (TBS + 0.05% Tween 20 + 0.1% BSA, pH 7.3) and, unless otherwise stated, steps were conducted at room temperature. Each wash involved a brief 20 s agitation, and all conjugation, blocking, and incubation steps were performed with gentle agitation over the entire stated time period. The general procedure for an ELISA assay was: wash 3×200 μL/well, conjugate with 2 μM of biotinylated compound for 2 h at room temperature (100 μL/well), wash 3×200 μL/well, blocked overnight in 5 wt% milk at 4°C, wash 3×200 μL/well with wash buffer, incubate with the titrated compound for 1 h, 3×200 μL/well, incubate with primary antibody (either or) for 1 h (100 μL/well), wash 3×200 μL/well, incubate with 1:2,000 anti-rabbit secondary antibody-horseradish peroxidase conjugate (Cell signaling Technologies, 7074S) for 1 h, 3×200 μL/well, develop with the Microwell Peroxidase Substrate System (2-C) (SeraCare, 5120-0047) using 100 μL/well for 1-40 minutes, and quench using 1 M H_2_SO_4_ (aq.) at 100 μL/well. For MrkA binding assays, biotinylated PCCs were conjugated to the well surface, recombinant MrkA with an N-terminal 6xHis-SUMO-tag (MyBiosource, MBS1248970) was titrated at the desired concentration, and the primary antibody was His-tag antibody, pAb, Rabbit (Genscript, A00174) at a 1:5,000 dilution. For epitope selectivity assays, biotinylated SynEps were conjugated, DNP-conjugated AR-PCCs were titrated at a desired concentration, and the primary antibody was anti-DNP antibody produced in Rabbit (Sigma-Aldrich, D9656) at a 1:8,000 dilution. The PCCs and SynEps used for plate-based ELISA assays had a peg5 linker between the N-terminus of the peptide and the tag, which was either biotin or DNP-modified lysine.

### Cell culture

Bacteria were grown overnight from glycerol stocks by inoculation into either minimal media containing 1% glycerol and 0.3% casamino acids (G-CAA) or Lysogeny Broth (LB) with shaking at 37°C. An immortalised mouse bone marrow derived macrophage cell line (iBMDM, a kind gift from Dr. Eicke Latz) was used in this study. Cells were maintained in RPMI containing 10% serum at 37°C in a humidified atmosphere with 5% CO_2_.

### Salt aggregation tests

Bacterial were cultured overnight in G-CAA broth, washed once in 0.02M phosphate buffer (0.01 M Na_2_HPO_4_, 0.01 M NaH_2_PO_4_, pH 8), and then resuspended to an O.D._600_ of 0.95. The bacterial solutions were arrayed onto a single glass slide in 10 μL spots, and then each spot was mixed with an equal volume of phosphate buffer containing (NH_4_)_2_SO_4_. The glass slide was gently agitated for 2 m. Images were recorded at 10 m and 30 m following agitation, and image acquisition of all the spots took less than 1 m. Measurements were conducted on both live bacteria and heat-killed bacteria that were treated for 10 minutes at 90 °C.

### Cell-based ELISAs

Detection of the binding of biotinylated PCCs to bacterial surface was performed as follows. 10^8^ bacteria from a culture grown overnight were used for each test. Bacteria were incubated with PBS containing 1% BSA (PBS-BSA) for 1h at 37°C, washed once with PBS, and then incubated with 50 µM of the biotinylated PCC in PBS-BSA for 1 h at 37°C. After washing thrice with PBS to remove unbound PCCs, bacteria were incubated with Streptavidin-HRP (1:1000) for 1 h at 37°C. Bacteria were washed thoroughly and the TMB reagent was added until visible coloration was observed. The reaction was quenched using 2N H_2_SO_4_ and absorbance was measured at 450 nm. The biotinylated PCCs used for cell-based ELISA assays had a peg5 linker between the N-terminus of the PCC and the biotin tag.

The protocol for detecting anti-DNP recruitment to bacterial cell surfaces is as follows. Bacteria were incubated with the desired concentration of DNP-conjugated PCCs in PBS-BSA for 1 h at 37°C. Residual PCCs were washed off by using PBS-BSA and bacteria were incubated with anti-DNP antibody (1:1000) for 1 h at 37°C. After washing, bacteria were incubated with HRP conjugated secondary antibody (Bio-Rad) at a dilution of 1:10,000 for 1 h at 37°C. Unbound antibody was washed off by using PBS-BSA and the cells were then developed by using TMB reagent. The reaction was quenched using 2N H_2_SO_4_ and absorbance was measured at 450 nm. Anti-MrkA antibody was procured from Biorbyt (orb51318) and used at a concentration of 5 μg/mL, and *K. pneumoniae* antiserum was obtained from abcam (ab20947) and used at a concentration of 5 μg/mL. The DNP-conjugated PCCs used for anti-DNP recruitment assays had a peg5-peg5 linker between the N-terminus of the PCC and the DNP-modified lysine.

### Opsonophagocytic killing (OPK) assays

*Klebsiella pneumoniae* BAA1705 was treated with DNP-PCCs followed by incubation with anti-DNP antibody, as described above in the Cell-Based ELISAs section. These opsonized bacteria were then used to infect BMDMs at a multiplicity of infection of 50 for 0.5 h at 37°C. Bacteria were then washed by using RPMI 1640 Media and BMDMs were left in cell culture medium containing gentamicin (100µg/ml). After 1 h and 24 h BMDMs were washed to remove gentamicin and intracellular bacteria were harvested by lysing BMDMs in RPMI media containing 0.2% Triton X 100. Bacterial CFUs were enumerated by plating onto LB agar. Bacterial counts at 1 h indicated the degree of opsonization while those at 24h served as a measure of microbicidal activity of macrophages. For these measurements, anti-MrkA antibody and *K*. pneumoniae antiserum were used at dilutions of 5 μg/mL each. The DNP-conjugated PCCs used for the phagocytosis and OPK assays had a peg5-peg5 linker between the N-terminus of the PCC and the DNP-modified lysine.

### Flow cytometry

Cytometry measurements were used to quantify AR-PCC-driven opsonization and opsonophagocytic killing of *K. pneumoniae* cells. For these measurements, *K. pneumoniae* cells were cultured in G-CAA medium overnight, washed, incubated with DNP-tagged AR-PCC at a desired concentration, washed, incubated with anti-DNP antibody conjugated with Alexafluor488, washed, then resuspended in PBS media. The samples were stored at 4 °C for 2 d before cytometry measurements were performed. The cytometer was calibrated by using Rainbow fluorescent beads (3.5 µm diameter) 3.5 m from BD (559123), which aided identification of single *K. pneumoniae* cells in subsequent measurements. Samples were excited with 488 nm light and the fluorescence emission at 530 nm was measured. Sample with no Alexaflour488 stain was used as fluorescent minus one (FMO) control for gating of DNP+ population. FACS data was analyzed on FlowJo v10 software, and the resulting histograms each include fluorescence data from >18,000 cells for strain BAA 2146 (Figure 4(B)) and >4,000 cells for strain BAA 1705 (Figure S10, SI). PCCs used for flow cytometry assays had a peg5-peg5 linker between the N-terminus of the PCC and the DNP tag.

## Acknowledgements

The acknowledgements come at the end of an article after the conclusions and before the notes and references.

