## Supplementary Information for "Antibody-Recruiting Protein-Catalyzed Capture Agents to Combat Antibiotic-Resistant Bacteria"

### SUPPORTING INFORMATION

**Table S1.** Predicted and average surface exposure, antigenicity and maximum homology for the sequence of MrkA.

| Residue # | Residue | Predicted quantities | | | | |
|---|---|---|---|---|---|---|
| | | Pred. Antigen. | Pred. Surf.-Exp. | Avg. Surf.-Exp. | Avg. Antigen. | Max Homol. (%) |
| 1 | M | 0 | 1 | 29 | 0 | 64 |
| 2 | K | 0 | 1 | 21 | 0 | 64 |
| 3 | K | 0 | 1 | 14 | 0 | 71 |
| 4 | V | 0 | 1 | 7 | 0 | 71 |
| 5 | L | 0 | 0 | 0 | 0 | 71 |
| 6 | L | 0 | 0 | 0 | 0 | 64 |
| 7 | S | 0 | 0 | 0 | 0 | 57 |
| 8 | A | 0 | 0 | 0 | 0 | 57 |
| 9 | A | 0 | 0 | 0 | 0 | 57 |
| 10 | M | 0 | 0 | 0 | 0 | 57 |
| 11 | A | 0 | 0 | 0 | 7 | 57 |
| 12 | T | 0 | 0 | 0 | 14 | 57 |
| 13 | A | 0 | 0 | 7 | 21 | 57 |
| 14 | F | 0 | 0 | 7 | 29 | 57 |
| 15 | F | 0 | 0 | 14 | 36 | 57 |
| 16 | G | 0 | 0 | 21 | 43 | 64 |
| 17 | M | 0 | 0 | 21 | 50 | 71 |
| 18 | T | 0 | 0 | 29 | 50 | 71 |
| 19 | A | 0 | 0 | 29 | 50 | 71 |
| 20 | A | 0 | 0 | 36 | 50 | 71 |
| 21 | H | 0 | 0 | 36 | 50 | 64 |
| 22 | A | 0 | 0 | 43 | 50 | 64 |
| 23 | A | 0 | 0 | 43 | 50 | 57 |
| 24 | D | 1 | 0 | 50 | 50 | 64 |
| 25 | T | 1 | 0 | 50 | 43 | 64 |
| 26 | T | 1 | 1 | 57 | 36 | 64 |
| 27 | V | 1 | 0 | 57 | 29 | 64 |
| 28 | G | 1 | 1 | 64 | 21 | 64 |
| 29 | G | 1 | 1 | 64 | 14 | 64 |
| 30 | G | 1 | 0 | 57 | 7 | 50 |
| 31 | Q | 0 | 1 | 64 | 0 | 64 |
| 32 | V | 0 | 0 | 57 | 0 | 64 |
| 33 | N | 0 | 1 | 64 | 0 | 64 |
| 34 | F | 0 | 0 | 64 | 0 | 57 |
| 35 | F | 0 | 1 | 71 | 7 | 57 |
| 36 | G | 0 | 0 | 71 | 14 | 57 |
| 37 | K | 0 | 1 | 79 | 21 | 64 |
| 38 | V | 0 | 0 | 79 | 29 | 64 |
| 39 | T | 0 | 1 | 86 | 36 | 64 |
| 40 | D | 0 | 1 | 86 | 43 | 57 |
| 41 | V | 0 | 1 | 79 | 50 | 57 |
| 42 | S | 0 | 1 | 79 | 50 | 57 |
| 43 | C | 0 | 0 | 71 | 50 | 64 |
| 44 | T | 0 | 1 | 79 | 50 | 57 |
| 45 | V | 0 | 0 | 71 | 50 | 57 |
| 46 | S | 0 | 1 | 79 | 50 | 57 |
| 47 | V | 0 | 1 | 79 | 50 | 71 |
| 48 | N | 1 | 1 | 71 | 57 | 64 |
| 49 | G | 1 | 1 | 71 | 50 | 64 |
| 50 | Q | 1 | 1 | 71 | 50 | 71 |
| 51 | G | 1 | 1 | 71 | 50 | 57 |
| 52 | S | 1 | 1 | 71 | 50 | 64 |
| 53 | D | 1 | 1 | 64 | 50 | 57 |
| 54 | A | 1 | 0 | 64 | 50 | 57 |
| 55 | N | 0 | 1 | 71 | 50 | 64 |
| 56 | V | 0 | 0 | 71 | 57 | 64 |
| 57 | Y | 0 | 1 | 79 | 64 | 64 |
| 58 | L | 0 | 0 | 79 | 71 | 64 |
| 59 | S | 0 | 1 | 86 | 79 | 71 |
| 60 | P | 0 | 1 | 86 | 86 | 71 |
| 61 | V | 1 | 0 | 86 | 93 | 64 |
| 62 | T | 0 | 1 | 93 | 93 | 64 |
| 63 | L | 1 | 1 | 93 | 100 | 71 |
| 64 | T | 1 | 1 | 93 | 100 | 71 |
| 65 | E | 1 | 1 | 93 | 100 | 71 |
| 66 | V | 1 | 0 | 93 | 93 | 64 |
| 67 | K | 1 | 1 | 93 | 86 | 57 |
| 68 | A | 1 | 1 | 93 | 79 | 57 |
| 69 | A | 1 | 1 | 86 | 71 | 57 |
| 70 | A | 1 | 1 | 86 | 64 | 57 |
| 71 | A | 1 | 1 | 79 | 57 | 64 |
| 72 | D | 1 | 1 | 79 | 50 | 64 |
| 73 | T | 1 | 1 | 79 | 43 | 64 |
| 74 | Y | 1 | 1 | 71 | 43 | 50 |
| 75 | L | 1 | 1 | 71 | 43 | 57 |
| 76 | K | 1 | 1 | 71 | 43 | 57 |
| 77 | P | 1 | 1 | 71 | 43 | 57 |
| 78 | K | 1 | 1 | 71 | 43 | 57 |
| 79 | S | 0 | 1 | 71 | 43 | 57 |
| 80 | F | 0 | 0 | 71 | 50 | 57 |
| 81 | T | 0 | 1 | 79 | 57 | 57 |
| 82 | I | 0 | 0 | 79 | 64 | 50 |
| 83 | D | 0 | 1 | 86 | 71 | 57 |
| 84 | V | 0 | 0 | 86 | 79 | 57 |
| 85 | S | 0 | 1 | 86 | 86 | 57 |
| 86 | N | 0 | 1 | 86 | 86 | 57 |
| 87 | C | 1 | 0 | 86 | 86 | 57 |
| 88 | Q | 1 | 1 | 86 | 79 | 57 |
| 89 | A | 1 | 1 | 86 | 71 | 57 |
| 90 | A | 1 | 1 | 79 | 64 | 57 |
| 91 | D | 1 | 1 | 71 | 57 | 71 |
| 92 | G | 1 | 1 | 64 | 50 | 57 |
| 93 | T | 1 | 1 | 64 | 43 | 57 |
| 94 | K | 1 | 1 | 57 | 36 | 64 |
| 95 | Q | 1 | 1 | 50 | 36 | 64 |
| 96 | D | 1 | 1 | 50 | 36 | 64 |
| 97 | D | 1 | 1 | 50 | 36 | 57 |
| 98 | V | 1 | 0 | 43 | 36 | 57 |
| 99 | S | 0 | 1 | 50 | 36 | 57 |
| 100 | K | 0 | 1 | 50 | 43 | 57 |
| 101 | L | 0 | 0 | 50 | 50 | 57 |
| 102 | G | 0 | 1 | 57 | 57 | 57 |
| 103 | V | 0 | 0 | 57 | 64 | 64 |
| 104 | N | 0 | 0 | 64 | 71 | 64 |
| 105 | W | 0 | 0 | 64 | 79 | 57 |
| 106 | T | 0 | 1 | 71 | 86 | 64 |
| 107 | G | 0 | 0 | 71 | 93 | 57 |
| 108 | G | 1 | 0 | 71 | 100 | 57 |
| 109 | N | 1 | 1 | 71 | 100 | 57 |
| 110 | L | 1 | 1 | 71 | 100 | 64 |
| 111 | L | 1 | 0 | 71 | 100 | 64 |
| 112 | A | 1 | 1 | 79 | 100 | 64 |
| 113 | G | 1 | 1 | 79 | 100 | 64 |
| 114 | A | 1 | 1 | 79 | 100 | 57 |
| 115 | T | 1 | 1 | 79 | 100 | 71 |
| 116 | S | 1 | 1 | 79 | 100 | 71 |
| 117 | K | 1 | 1 | 71 | 100 | 57 |
| 118 | Q | 1 | 0 | 71 | 93 | 64 |
| 119 | Q | 1 | 1 | 71 | 86 | 71 |
| 120 | G | 1 | 1 | 64 | 79 | 71 |
| 121 | Y | 1 | 0 | 57 | 71 | 57 |
| 122 | L | 1 | 0 | 57 | 64 | 57 |
| 123 | A | 1 | 1 | 57 | 57 | 57 |
| 124 | N | 1 | 1 | 50 | 50 | 57 |
| 125 | T | 1 | 1 | 43 | 43 | 64 |
| 126 | E | 1 | 1 | 43 | 36 | 57 |
| 127 | A | 1 | 1 | 43 | 36 | 57 |
| 128 | S | 1 | 1 | 43 | 36 | 57 |
| 129 | G | 1 | 1 | 43 | 36 | 57 |
| 130 | A | 1 | 0 | 43 | 36 | 57 |
| 131 | Q | 0 | 1 | 50 | 36 | 64 |
| 132 | N | 0 | 0 | 50 | 43 | 64 |
| 133 | I | 0 | 0 | 57 | 50 | 64 |
| 134 | Q | 0 | 0 | 64 | 57 | 64 |
| 135 | L | 0 | 0 | 71 | 64 | 71 |
| 136 | V | 0 | 0 | 71 | 71 | 57 |
| 137 | L | 0 | 0 | 79 | 79 | 57 |
| 138 | S | 0 | 0 | 86 | 86 | 57 |
| 139 | T | 0 | 1 | 93 | 93 | 64 |
| 140 | D | 1 | 1 | 93 | 100 | 64 |
| 141 | N | 1 | 1 | 93 | 100 | 64 |
| 142 | A | 1 | 1 | 93 | 100 | 57 |
| 143 | T | 1 | 1 | 93 | 100 | 64 |
| 144 | A | 1 | 1 | 93 | 100 | 71 |
| 145 | L | 1 | 1 | 93 | 100 | 64 |
| 146 | T | 1 | 1 | 93 | 100 | 64 |
| 147 | N | 1 | 1 | 93 | 100 | 57 |
| 148 | K | 1 | 1 | 93 | 100 | 57 |
| 149 | I | 1 | 0 | 93 | 100 | 57 |
| 150 | I | 1 | 1 | 100 | 100 | 57 |
| 151 | P | 1 | 1 | 100 | 100 | 64 |
| 152 | G | 1 | 1 | 100 | 100 | 64 |
| 153 | D | 1 | 1 | 100 | 100 | 64 |
| 154 | S | 1 | 1 | 100 | 100 | 57 |
| 155 | T | 1 | 1 | 100 | 100 | 64 |
| 156 | Q | 1 | 1 | 100 | 100 | 64 |
| 157 | P | 1 | 1 | 93 | 100 | 64 |
| 158 | K | 1 | 1 | 93 | 93 | 57 |
| 159 | A | 1 | 1 | 86 | 86 | 64 |
| 160 | K | 1 | 1 | 86 | 79 | 64 |
| 161 | G | 1 | 1 | 79 | 71 | 64 |
| 162 | D | 1 | 1 | 71 | 64 | 64 |
| 163 | A | 1 | 1 | 64 | 57 | 64 |
| 164 | S | 1 | 1 | 57 | 50 | 71 |
| 165 | A | 1 | 1 | 50 | 43 | 71 |
| 166 | V | 1 | 1 | 43 | 36 | 64 |
| 167 | A | 1 | 1 | 36 | 29 | 64 |
| 168 | D | 1 | 1 | 36 | 21 | 64 |
| 169 | G | 1 | 1 | 36 | 14 | 57 |
| 170 | A | 1 | 0 | 36 | 7 | 64 |
| 171 | R | 0 | 1 | 43 | 7 | 57 |
| 172 | F | 0 | 0 | 43 | 14 | 57 |
| 173 | T | 0 | 1 | 43 | 14 | 64 |
| 174 | Y | 0 | 0 | 43 | 14 | 64 |
| 175 | Y | 0 | 0 | 50 | 14 | 64 |
| 176 | V | 0 | 0 | 57 | 14 | 64 |
| 177 | G | 0 | 0 | 64 | 14 | 64 |
| 178 | Y | 0 | 0 | 64 | 14 | 71 |
| 179 | A | 0 | 0 | 71 | 14 | 71 |
| 180 | T | 0 | 0 | 71 | 14 | 71 |
| 181 | S | 0 | 1 | 79 | 14 | 64 |
| 182 | A | 0 | 1 | 71 | 14 | 57 |
| 183 | P | 0 | 1 | 71 | 14 | 57 |
| 184 | T | 1 | 1 | 64 | 14 | 57 |
| 185 | T | 1 | 1 | 57 | 7 | 57 |
| 186 | V | 0 | 0 | 50 | 0 | 50 |
| 187 | T | 0 | 1 | 57 | 0 | 57 |
| 188 | T | 0 | 1 | 50 | 0 | 50 |
| 189 | G | 0 | 1 | 50 | 0 | 50 |
| 190 | V | 0 | 1 | — | — | — |
| 191 | V | 0 | 0 | — | — | — |
| 192 | N | 0 | 1 | — | — | — |
| 193 | S | 0 | 0 | — | — | — |
| 194 | Y | 0 | 1 | — | — | — |
| 195 | A | 0 | 0 | — | — | — |
| 196 | T | 0 | 1 | — | — | — |
| 197 | Y | 0 | 0 | — | — | — |
| 198 | E | 0 | 0 | — | — | — |
| 199 | I | 0 | 0 | — | — | — |
| 200 | T | 0 | 1 | — | — | — |
| 201 | Y | 0 | 0 | — | — | — |
| 202 | Q | 0 | 1 | — | — | — |

**Table S2.** Isocratic points and relative affinities of macrocyclic analogues of lead MrkA ligand cy(LLFFF).

| ELISA Assay # | Compound # | N-terminal tag | Cycle sequence | Isocratic point (%) | Relative affinity | [MrkA] for affinity test | Modification |
|---|---|---|---|---|---|---|---|
| ELISA 1 | Ref. compound | Bio-Peg10-peg10- | cy(X-LLFFF-Z) | 74.53 | 1.00 | N/A | Reference |
| ELISA 2 | Ref. compound | Bio-peg10- | cy(X-LLFFF-Z) | 75.88 | 1.00 | 25 nM | Reference |
| | 2 | Bio-peg10- | cy(X-ALFFF-Z) | 65.55 | 0.53 | 25 nM | L-to-A substitution |
| | 3 | Bio-peg10- | cy(X-LAFFF-Z) | 66.00 | 0.26 | 25 nM | L-to-A substitution |
| | 4 | Bio-peg10- | cy(X-LLAFF-Z) | 70.97 | 0.59 | 25 nM | F-to-A substitution |
| | 5 | Bio-peg10- | cy(X-LLFAF-Z) | 66.00 | 0.10 | 25 nM | F-to-A substitution |
| | 6 | Bio-peg10- | cy(Z-LLFFA-X) | 64.43 | 0.04 | 25 nM | F-to-A substitution |
| ELISA 3 | Ref. compound | Bio-peg10- | cy(X-LLFFF-Z) | 75.88 | 1.00 | 25 nM | Reference |
| | 7 | Bio-peg10- | cy(X-AAFFF-Z) | 54.65 | 0.15 | 50 nM | LL-to-AA substitution |
| | 8 | Bio-peg10- | cy(X-LFFF-Z) | 61.84 | 0.46 | 50 nM | L removal |
| | 10 | Bio-peg10- | cy(X-LLFFF-(Z3)) | 76.89 | 0.89 | 50 nM | Z-to-Z3 substitution (remove one methylene group) |
| | 11 | Bio-peg10- | cy(X-LLFF4-Z) | 75.53 | 0.88 | 50 nM | F-to-4 substitution |
| ELISA 4 | Ref. compound | Bio-peg5- | cy(X-LLFFF-Z) | 78.85 | 1.00 | 100 nM | Reference |
| | 13 | Bio-peg5- | cy(X-TTFFF-Z) | 53.40 | 0.04 | 100 nM | LL-to-TT substitution |
| ELISA 5 | Ref. compound | Bio-peg10- | cy(X-LLFFF-Z) | 75.88 | 1.00 | 25 nM | Reference |
| | 16 | Bio-peg10-R3-peg10- | cy(X-LLFFF-Z) | 45.02 | 0.90 | 25 nM | Add -Peg10-R3 tag |
| | 17 | Bio-peg10-R6-peg10- | cy(X-LLFFF-Z) | 39.82 | 1.22 | 25 nM | Add -Peg10-R6 tag |
| | 18 | Bio-peg10-R9-peg10- | cy(X-LLFFF-Z) | 37.03 | 0.58 | 25 nM | Add -Peg10-R9 tag |
| ELISA 6 | Ref. compound | Bio-peg10- | cy(X-LLFFF-Z) | 75.88 | 1.00 | 25 nM | Reference |
| | 19 | Bio-peg10- | cy(X-HLFFF-Z) | 47.52 | 0.81 | 25 nM | Cationic residue substitution |
| | 20 | Bio-peg10- | cy(X-DLFFF-Z) | 66.33 | 0.21 | 25 nM | Anionic residue substitution |
| | 21 | Bio-peg10- | cy(X-LLFF-Z) | 64.99 | 0.14 | 25 nM | Residue removal |
| | 22 | Bio-peg10- | cy(X-LLF-Z) | 56.45 | 0.09 | 25 nM | Double residue removal |
| | 23 | Bio-peg10- | cy(X-FFF-Z) | 60.27 | 0.14 | 25 nM | Double residue removal |

*cy() indicates cyclization, X = propargylglycine, Z = Azidolysine, Z3 = azidonorvaline, 4 = 4-fluoro-phenylalanine Bio = biotin, R*i* = polyarginine tag with *i* arginine residues, peg*i* = polyethylene glycol chain of length *i*. All amino acids represented by single-letter amino acid codes.

**MrkA SynEp 1 - Biotin-peg5-FFGKVT(V->Z)SCTVSV**

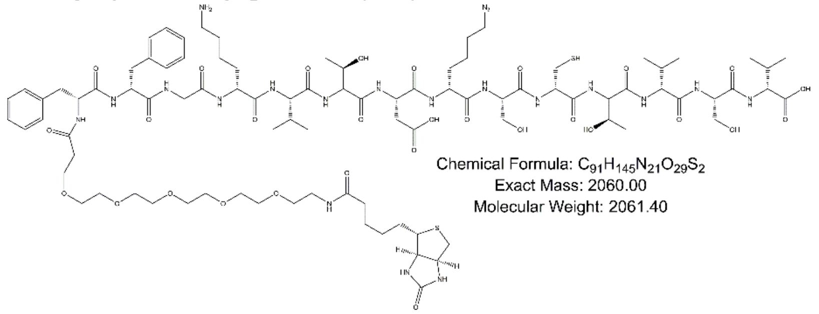


**MrkA SynEp 2 - Biotin-peg5-TEVKAA(A->Z)ADTYLKP**

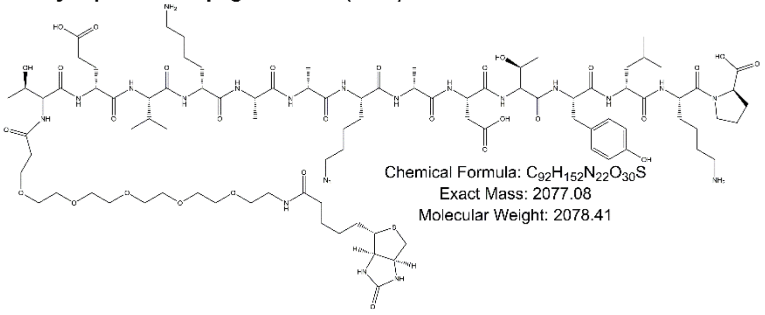


**MrkA SynEp 3 - Biotin-peg5-ATSKQQGY(L->Z)ANTEA**

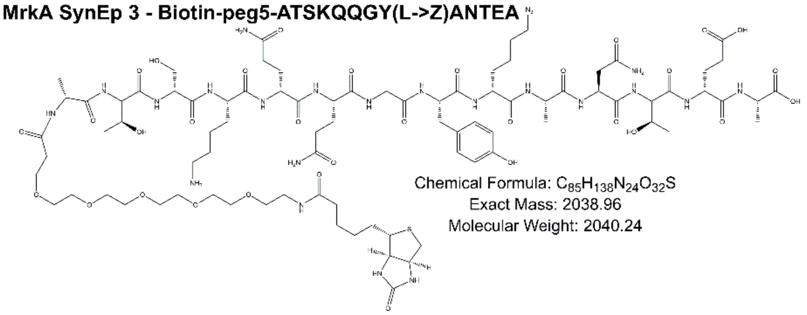


**MrkA SynEp 4 - Biotin-peg5-STQPK(A->Z)KGDASAVA**

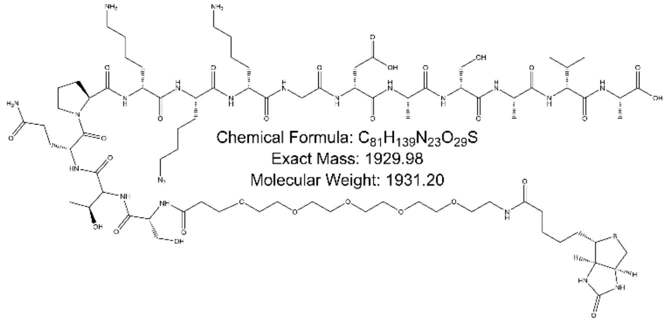


**Figure S1.** Molecular structures and parameters for synthetic epitopes used for PCC combinatorial screens in this study.

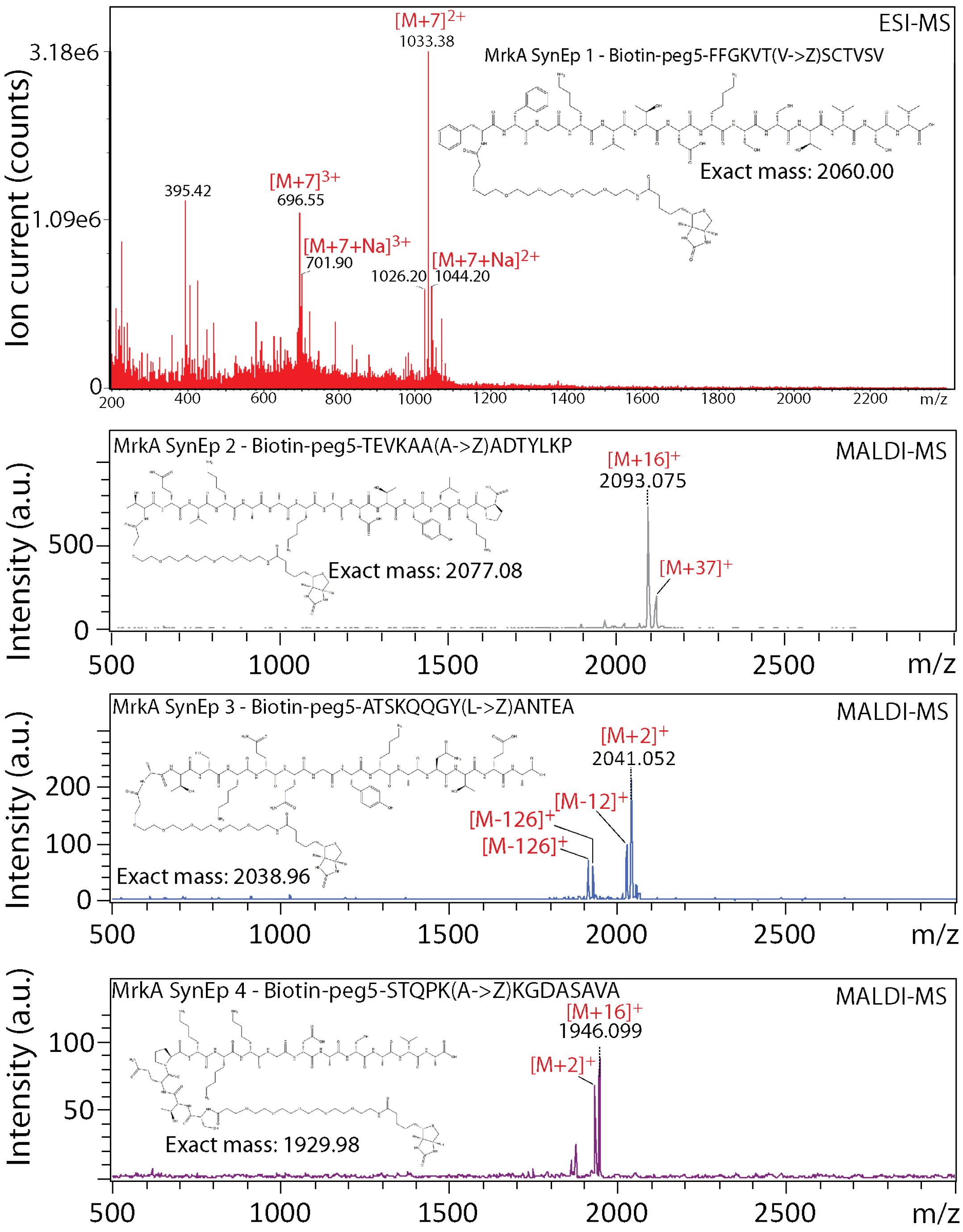


**Figure S2.** Representative mass spectroscopy characterization data for synthetic epitopes (SynEps) used for PCC combinatorial screening. Synthetic epitope 1 was purified by using a Waters Autopurification system that employs electrospray ionization (ESI) to detect and purify compounds, while SynEps 2-4 were purified by HPLC and then matrix-assisted laser desorption ionization spectroscopy. Multiple HPLC runs were used to purify each compound, and the mass spectroscopy data shown here are acquired from a single run or fraction that was subsequently pooled. Masses with +16 likely arise from oxidation of the SynEp.

(A)

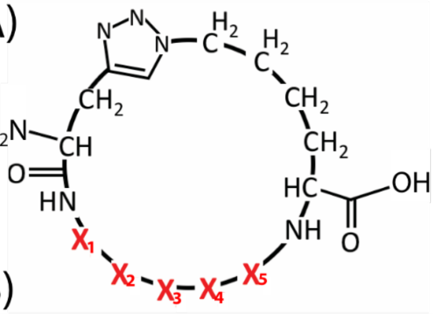


(B)

| Hit # | Protein | HPraGly | X1 | X2 | X3 | X4 | X5 | Az4 |
|---|---|---|---|---|---|---|---|---|
| 1 | MrkA | | F | Y | T | K | G | |
| 2 | MrkA | | E | Y | E | G | K | |
| 3 | MrkA | | P | W | N | K | G | |
| 4.1 | MrkA | | S | W | T | G | E | |
| 4.2 | MrkA | | R | H | P | G | E | |
| 5 | MrkA | | N | R | T | G | P | |
| 6 | MrkA | | P | R | E | G | P | |
| 7 | MrkA | | S | N | F | G | P | |
| 8 | MrkA | | E | K | T | P | G | |
| 9 | MrkA | | K | G | F | P | G | |
| 10 | MrkA | | G | G | F | N | A | |
| 11 | MrkA | | N | G | P | V | H | |
| 12 | MrkA | | W | Y | K | G | P | |
| 13 | MrkA | | W | D | Y | K | G | |
| 14 | MrkA | | Y | R | H | L | G | |
| 15 | MrkA | | G | V | H | R | L | |
| 16 | MrkA | | G | V | V | E | K | |
| 17 | MrkA | | G | L | T | H | A | |
| 18 | MrkA | | S | L | G | L | T | |
| 19 | MrkA | | L | L | F | F | F | |
| 20 | MrkA | | T | T | F | F | F | |
| 21 | MrkA | | K | P | A | G | | |
| 22 | MrkA | | A | K | P | E | P | |
| 23 | MrkA | | E | W | V | S | A | |
| 24 | MrkA | | E | F | S | G | V | |
| 25 | MrkA | | D | G | T | A | L | |
| 26.1 | MrkA | | V | V | N | L | P | |
| 26.2 | MrkA | | T | P | N | L | P | |
| 26.3 | MrkA | | R | P | E | G | P | |

**Figure S3. (A)** Chemical structure of macrocyclic peptide ligands to MrkA, in which each red $X_i$ represents one of 17 common amino acids (excluding methionine, cysteine, and isoleucine). **(B)** Table with the sequences of the 26 different macrocyclic peptide ligands from the in situ click screen of four target epitopes on MrkA.

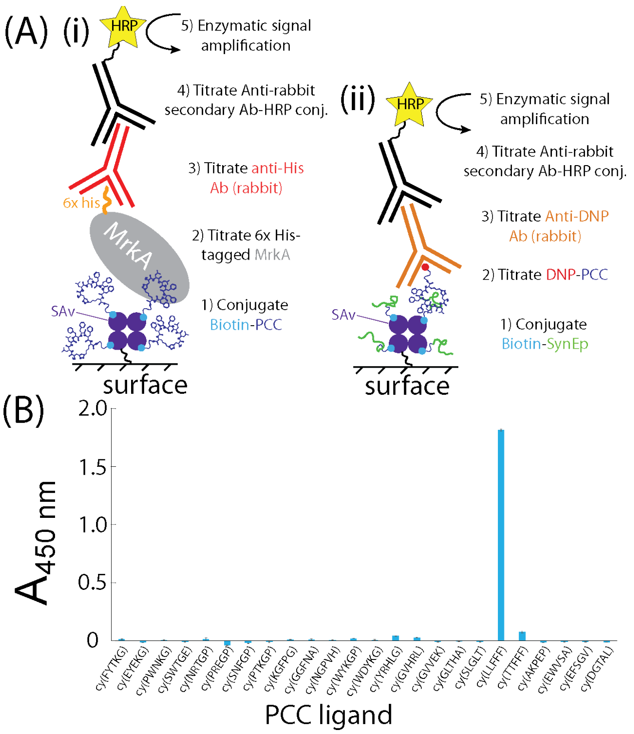


**Figure S4. (A)** Schematic diagrams of the sandwich ELISA assay formats used to test the binding affinities of PCC ligands to (i) full-length MrkA and (ii) SynEps. The relative sizes of the molecules in these illustrations are not to scale. **(B)** Bar graph showing the absorbances obtained from an enzyme-linked Immunoassay (ELISA) that tests the ability of immobilized PCC peptides to capture full-length MrkA at 100 nM from aqueous solution. The strong absorbance associated with immobilized *cy(LLFFF)* demonstrates strong binding of *cy(LLFFF)* to MrkA. Moderate binding is also observed for *cy(TTFFF)*, *cy(YRHLG)* and *cy(GVHRL)*. Measurements were done in duplicate and error bars indicate the span of the data. Intensities in (B) are shown after subtraction of a dummy ligand Biotin-peg5-cy(HNGPT) background.

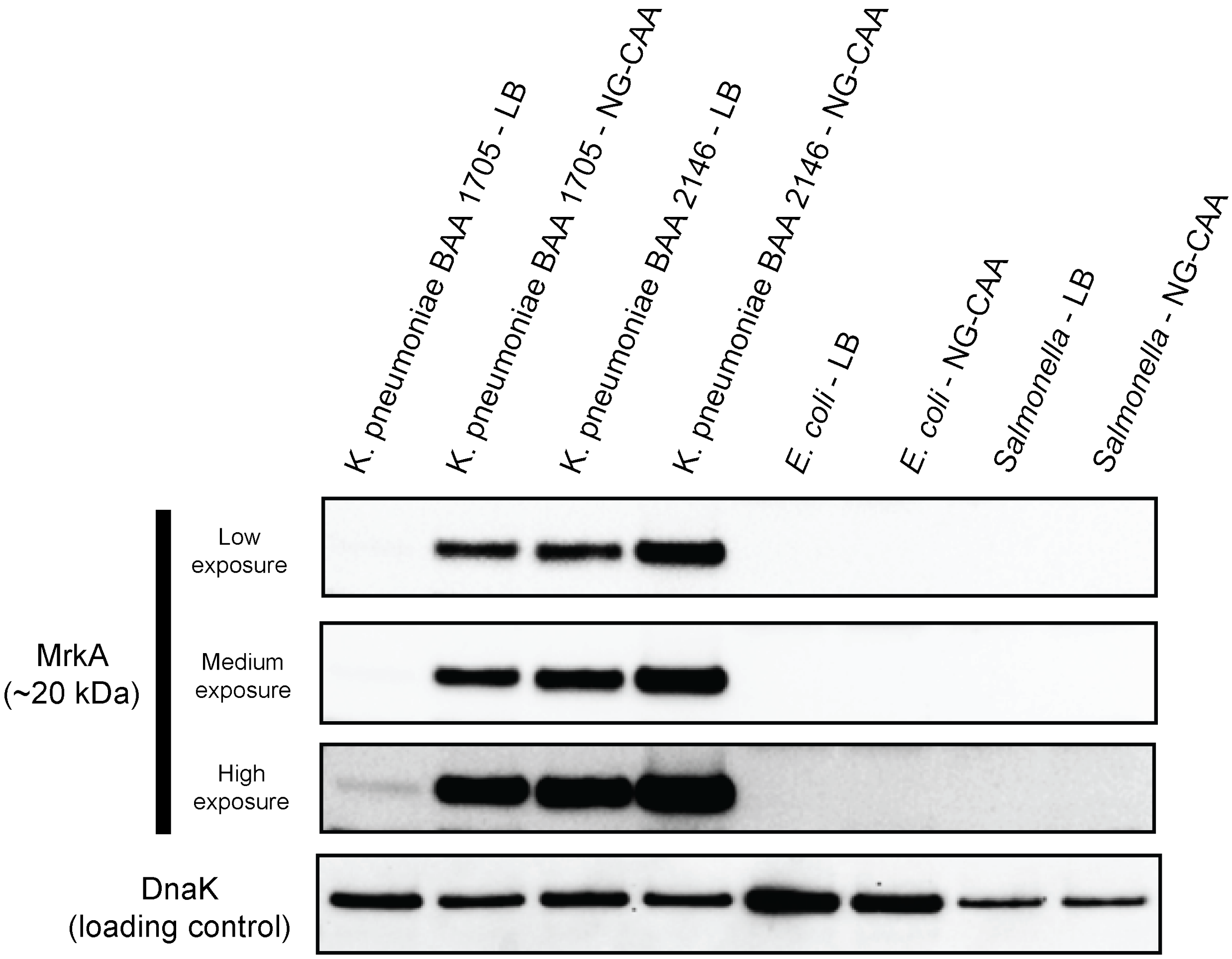


**Figure S5.** Western blot to detect MrkA protein in *Klebsiella pneumoniae*, *Escherichia coli* and *Salmonella typhimurium* grown in either LB (LB) or Glycerol Casamino acid solutions. *K. pneumoniae* BAA1705 grown in LB shows very fain signals associated with MrkA protein, while substantially higher signals associated with MrkA are observed for the same cells grown in NG-CAA. MrkA was highly expressed in *K. pneumoniae* BAA 2146 in either LB or GCAA. No detectable MrkA was observed in either *E. coli* or *S. typhimurium* under any growth conditions tested.

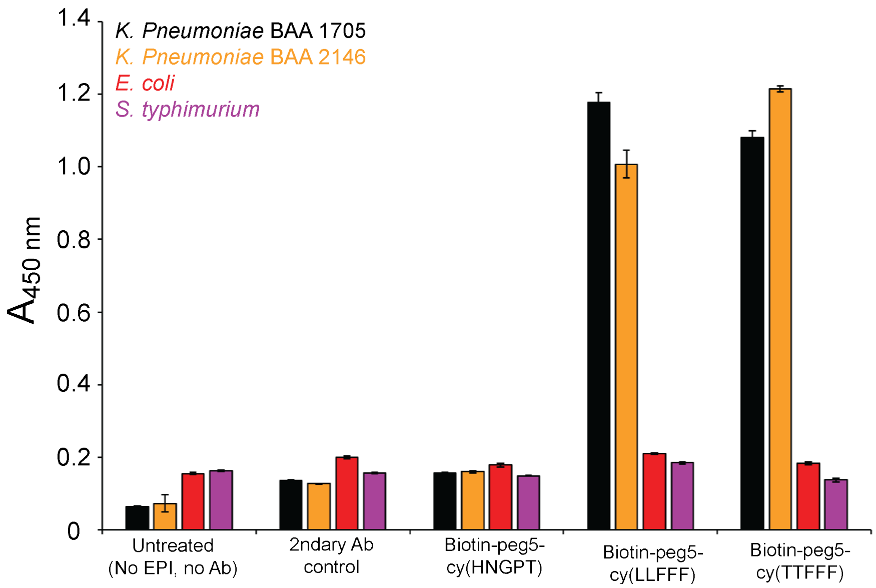


**Figure S6.** Plots of 450 nm absorbance for various bacterial cells opsonized without or with biotinylated AR-PCCs. Samples that were untreated or treated with secondary antibodies (Ab) show absorbances ranging from 0.05-0.2. Similar absorbances are observed from samples opsonized with the non-MrkA-binding biotin-peg5-cy(HNGPT) + 2ndary Ab, which establish little-to-no opsonization by this AR-PCC. By comparison, both biotin-peg5-cy(LLFFF) and biotin-peg5-cy(TTFFF) show strong absorbances from both *K. pneumoniae* strains and much smaller signals from *E. coli* or *S. typhimurium*. This establishes the selective opsonization of *K. pneumoniae* by lead AR-PCC ligands. The secondary antibody was streptavidin-horseradish peroxidase conjugate and all AR-PCCs were used at 50 μM concentrations.

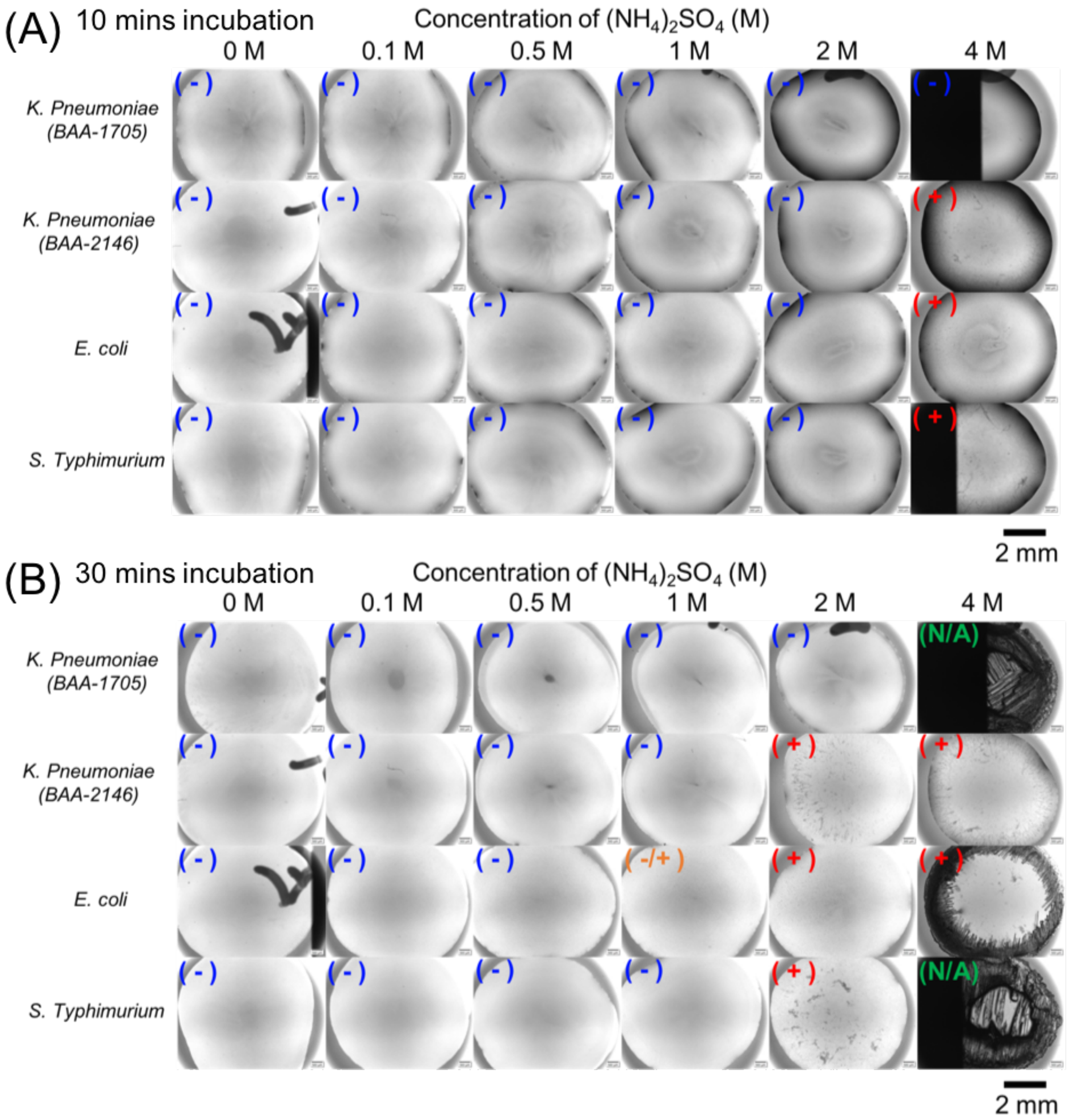


**Figure S7.** Images of droplets of phosphate buffered solution containing live bacteria and various concentrations of $(NH_4)_2SO_4$ at 10 m or 30 m following mixing. Aggregation in each droplet was assessed by visual inspection and is indicated as non-aggregated "( - )", low levels of aggregation "( -/+ )", and extensive aggregation "( + )". All droplets were on a single glass slide and image acquisition of all droplets took less than a minute for each 10 m and 30 m time point. Crystallization was observed at 30 m times, indicated by "N/A", that precluded assessment of aggregation. These results show that *K. pneumoniae* strains aggregate at similar or higher salt concentrations than *E. coli* and *S. typhimurium*, indicating similar or lower cell surface hydrophobicity for these strains of *K. pneumoniae* versus *E. coli* and *S. typhimurium*.

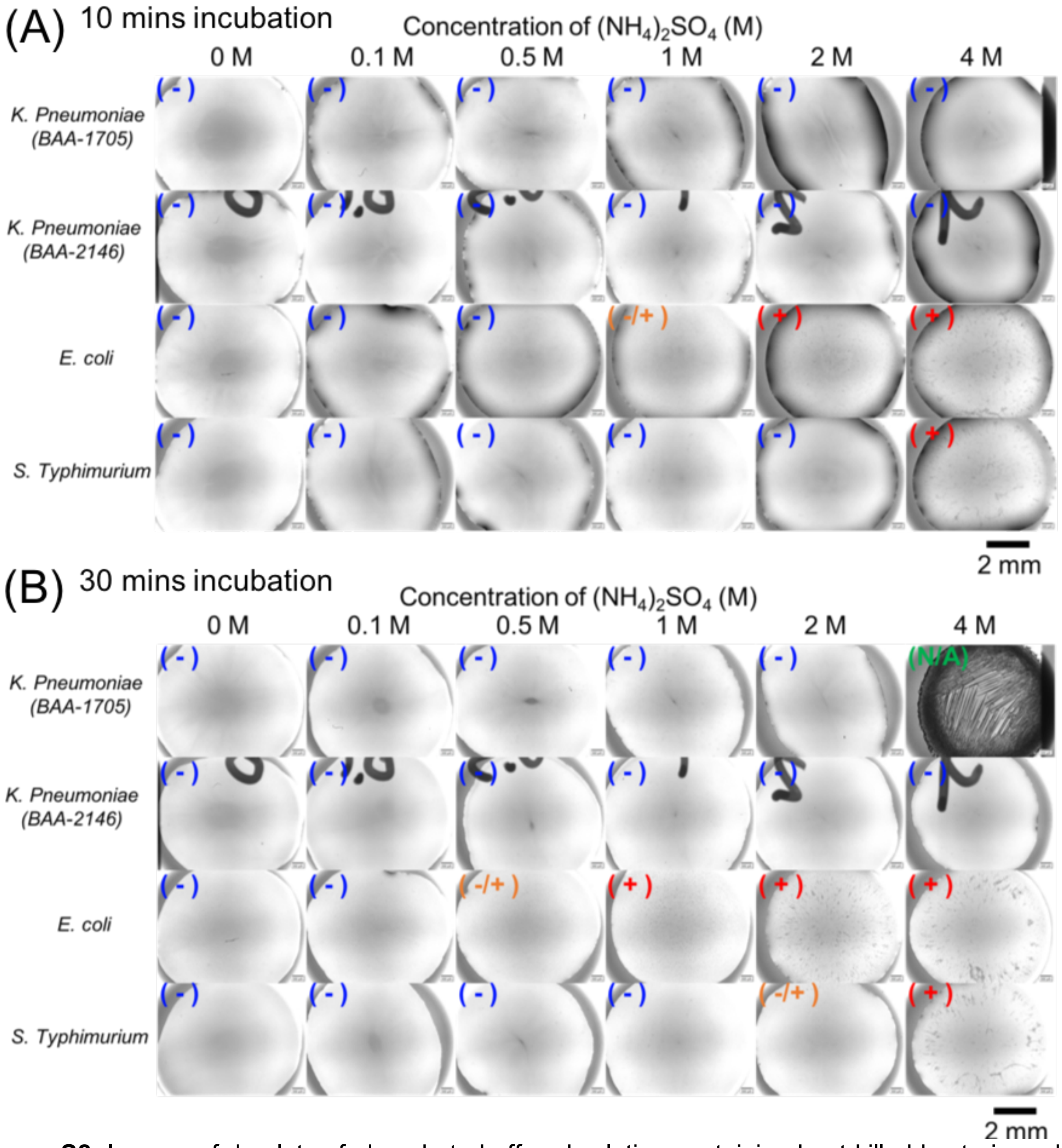


**Figure S8.** Images of droplets of phosphate buffered solution containing heat-killed bacteria and various concentrations of $(NH_4)_2SO_4$ at 10 m or 30 m following mixing. Aggregation in each droplet was assessed by visual inspection and is indicated as non-aggregated "( - )", low levels of aggregation "( -/+ )", and extensive aggregation "( + )". All droplets were on a single glass slide and image acquisition of all droplets took less than a minute for each 10 m and 30 m time point. Crystallization was observed at 30 m times, indicated by "N/A", that precluded assessment of aggregation. These results corroborate the similar cell surface hydrophobicities of *K. pneumoniae* strains used here and *E. coli* and *S. Typhimurium*, as indicated by salt aggregation tests on live bacteria.

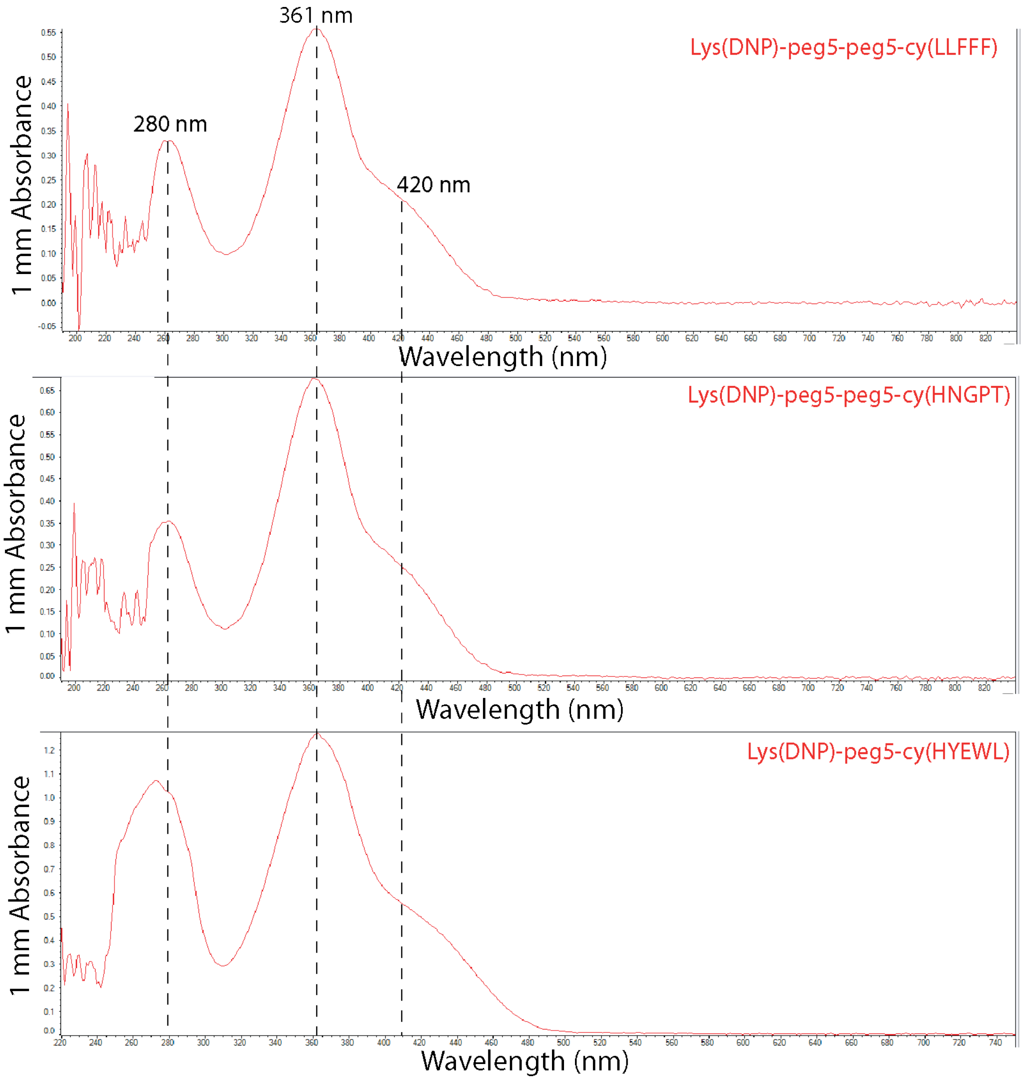


**Figure S9.** UV-visible absorbance spectra for DNP-conjugated compounds, including cy(LLFFF), cy(HNGPT) and cy(HYEWL), solubilized in dimethylsulfoxide, all of which show characteristic absorbance peak at 361 nm and shoulder at 420 nm that are associated with DNP. An absorbance peak at 280 nm is also present in each compound, which arises from amino acid residues of these compounds. DNP was conjugated to the macrocyclic peptide ligand via a C-terminal DNP-modified lysine residue, separated by either a peg5 or peg5-peg5 linker.

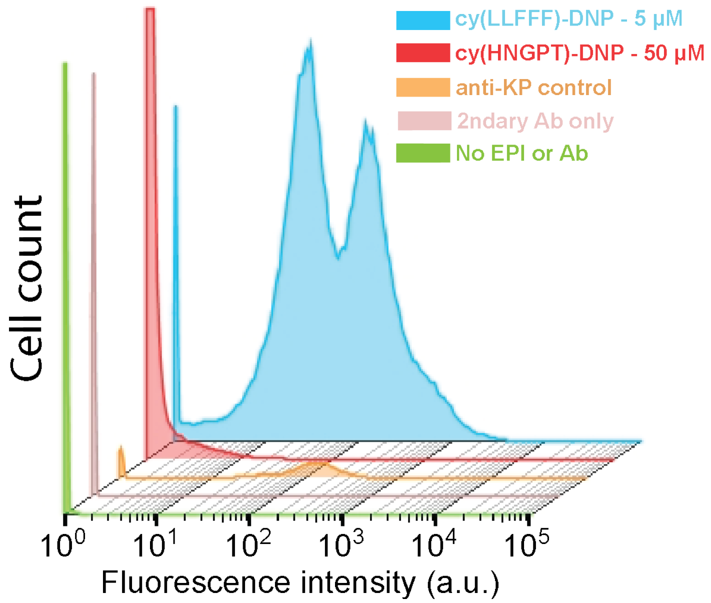


**Figure S10.** Plot of the fluorescence intensity versus cell count for *K. pneumoniae* (strain BAA 1705) opsonized by DNP-conjugated AR-PCC ligands and fluorescently-tagged anti-DNP antibodies. Cells exposed to cy(LLFFF) at 5 μM (blue) concentrations showed much higher fluorescence than cells incubated with 50 μM of the control cy(HGNPT)-DNP conjugate plus fluorescent anti-DNP (red). Cells exposed only to only *Klebsiella pneumoniae* antiserum (orange), secondary antibody (i.e., fluorescently tagged anti-DNP, pink), or neither AR-PCCs or antibodies (green) showed the least fluorescence. Cytometry measurements were gated for single cells and each histogram comprises >4,000 cells.

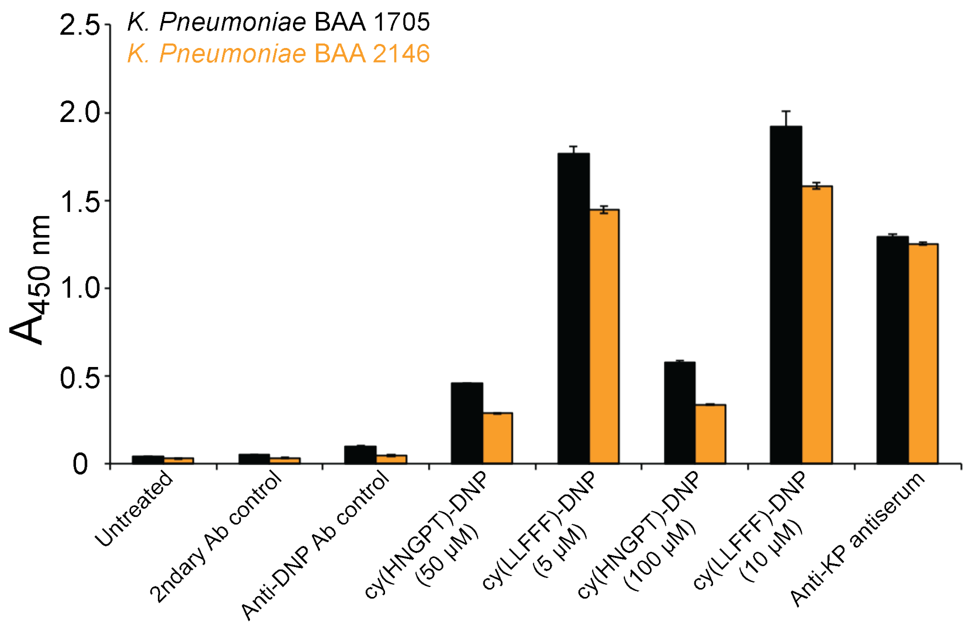


**Figure S11.** Plot of 450 nm absorbances obtained from a cell-based ELISA assay conducted on two strains of *K. pneumoniae* cells, BAA 1705 (black) and BAA 2146 (orange), that were either untreated or opsonized by AR-PCCs and/or antibodies. Very low absorbances were observed for *K. pneumoniae* cells that were untreated, or treated either with secondary antibody or anti-DNP controls. By comparison, much larger signal is observed for *K. pneumoniae* opsonized with cy(LLFFF) plus anti-DNP at either 5 μM or 10 μM AR-PCC concentrations (anti-DNP concentration fixed). This establishes AR-PCC-driven opsonization by cy(LLFFF)-DNP. Much smaller signals were observed from *K. pneumoniae* cells opsonized with a dummy AR-PCC ligand cy(HNGPT)-DNP at a much greater concentrations of 50 and 100 μM. This indicates opsonization by cy(LLFFF)-DNP is specific and is driven by the ligand cy(LLFFF), rather than the Lys(DNP) moiety.

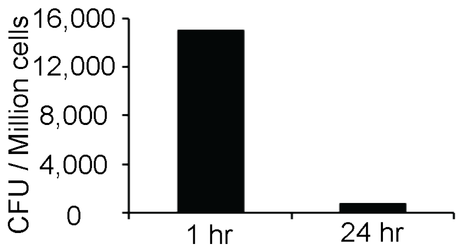


**Figure S12.** Plot of CFU counts per million cells obtained by plating and growing macrophage lysate after exposure of macrophages to *K. pneumoniae* bacteria for either 1 or 24 h. Larger cell counts (~15,000 CFU) were observed in samples prepared from macrophages harvested at 1 h, while very low counts (~1,000 CFU) were detected in samples from macrophages harvested at 24 h. This demonstrates that after 2 h, bacteria remain viable inside the phagosome, while after 24 h most bacteria were rendered inviable.

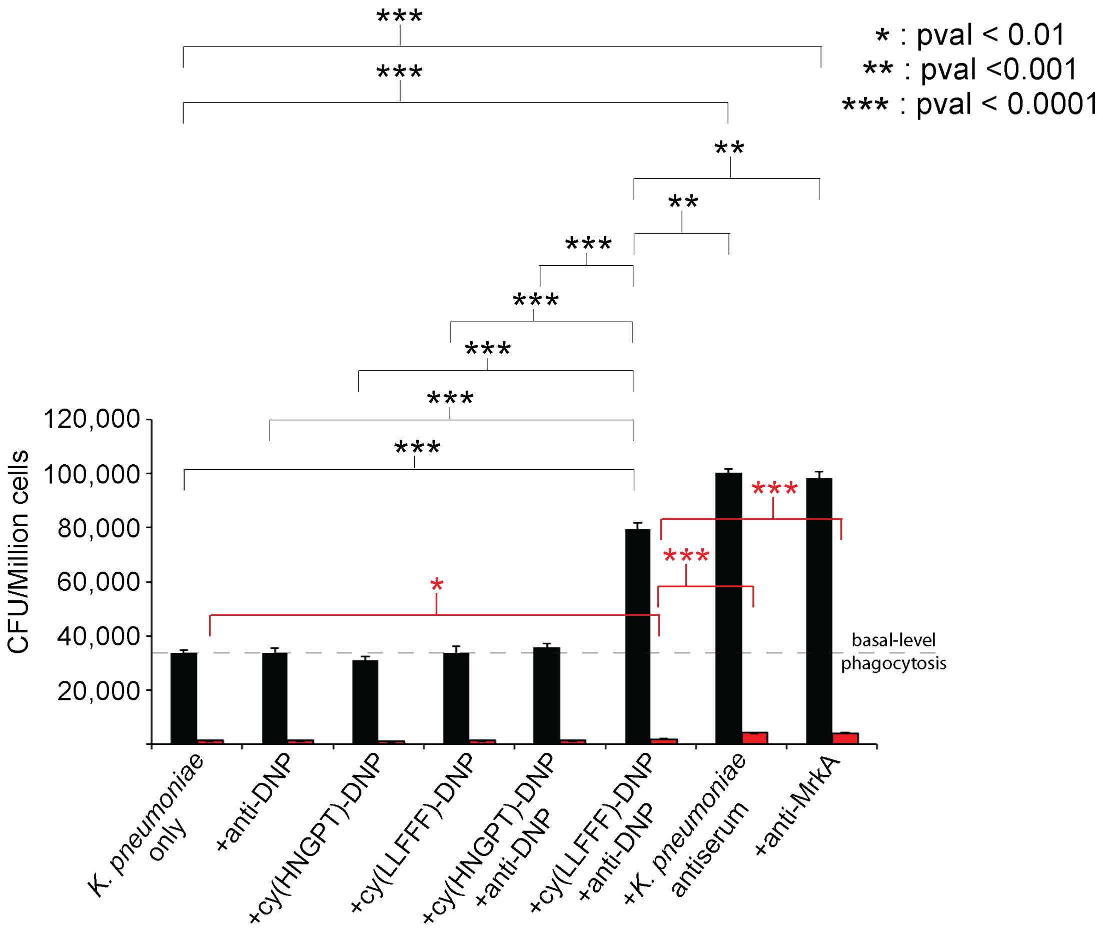


**Figure S13.** Statistical significance of the phagocytosis and opsonophagocytic killing (OPK) assays performed on *K. pneumoniae* without or with exposure to various compounds and antibodies, as shown in Figure 5 of the main text. Asterisks indicate the upper limit of the p-value for the corresponding t-tests (see legend) conducted on the colony forming unit (CFU) counts between two samples. Black and red brackets indicate comparisons of sample counts at 1 h and 24 h, respectively. For example, the topmost black bracket indicates a comparison between "K. pneumoniae only" and "+anti-MrkA" samples at 1 h, the three asterisks "***"indicate a p value of <0.001 for this t-test.

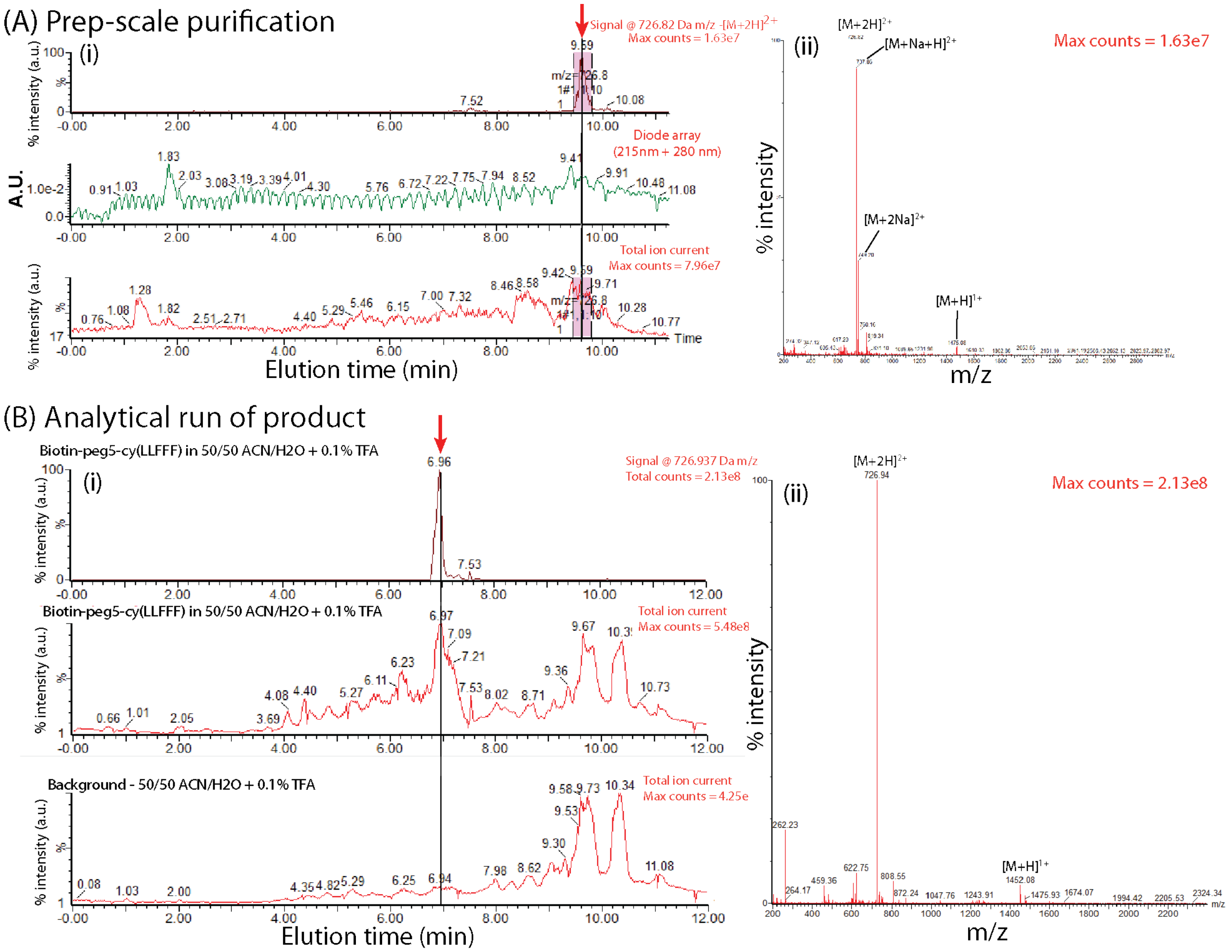


**Figure S14. (A)** Example purification and purity evaluation of the compound cy(LLFFF) conjugated with a peg5-biotin moiety. (i) HPLC-MS chromatograms of the total ion current (lower) and ion current at 726.82 +/- 2 Da (top), which is the $[M+2H]^{2+}$ mass of the desired product, and the absorbance (A280 + A215) (middle). The corresponding mass spectrum recorded at the elution time indicated by the red arrow in (i), which shows peaks associated with the $[M+H]^{+}$, $[M+2H]^{2+}$, $[M+Na+H]^{2+}$, $[M+2Na]^{2+}$. (B) (i) Example HPLC-MS chromatograms (top two) of the product purified in (A) in DMSO at 10 mM alongside a run on the same column but with 50/50 ACN/$H_2O$ without compound (lower). Shown in (ii) is a mass spectrum obtained at the elution time indicated in the red arrow in (i) which shows peaks associated with the $[M+H]^{+}$ and $[M+2H]^{2+}$.
